# Identification of novel Kv1.3 channel-interacting proteins using proximity labelling in T-cells

**DOI:** 10.1101/2025.01.16.633279

**Authors:** Dilpreet Kour, Christine A. Bowen, Upasna Srivastava, Hai M. Nguyen, Rashmi Kumari, Prateek Kumar, Amanda D. Brandelli, Sara Bitarafan, Brendan R Tobin, Levi Wood, Nicholas T. Seyfried, Heike Wulff, Srikant Rangaraju

**Affiliations:** Department of Neurology, School of Medicine, Yale University, New Haven (CT), USA; Center for Neurodegenerative Diseases, Emory University, Atlanta (GA), USA; Department of Biochemistry, Emory University, Atlanta (GA), USA; Department of Pharmacology, University of California – Davis, Davis (CA), USA; Parker H. Petit Institute for Bioengineering, Georgia Institute of Technology, Atlanta (GA), USA; George W. Woodruff School of Mechanical Engineering, Georgia Institute of Technology, Atlanta (GA), USA; School of Chemical and Biomolecular Engineering, Georgia Institute of Technology, Atlanta (GA), USA; Wallace H. Coulter Department of Biomedical Engineering, Georgia Institute of Technology, Atlanta (GA), USA

**Keywords:** Potassium channel, T cell, proteomics, autoimmune disease, interactions, proximity labeling

## Abstract

Potassium channels regulate membrane potential, calcium flux, cellular activation and effector functions of adaptive and innate immune cells. The voltage-activated Kv1.3 channel is an important regulator of T cell-mediated autoimmunity and microglia-mediated neuroinflammation. Kv1.3 channels, via protein-protein interactions, are localized with key immune proteins and pathways, enabling functional coupling between K+ efflux and immune mechanisms. To gain insights into proteins and pathways that interact with Kv1.3 channels, we applied a proximity-labeling proteomics approach to characterize protein interactors of the Kv1.3 channel in activated T-cells. Biotin ligase TurboID was fused to either N or C termini of Kv1.3, stably expressed in Jurkat T cells and biotinylated proteins in proximity to Kv1.3 were enriched and quantified by mass spectrometry. We identified over 1,800 Kv1.3 interactors including known interactors (beta-integrins, Stat1) although majority were novel. We found that the N-terminus of Kv1.3 preferentially interacts with protein synthesis and protein trafficking machinery, while the C-terminus interacts with immune signaling and cell junction proteins. T-cell Kv1.3 interactors included 335 cell surface, T-cell receptor complex, mitochondrial, calcium and cytokine-mediated signaling pathway and lymphocyte migration proteins. 178 Kv1.3 interactors in T-cells also represent genetic risk factors of T cell-mediated autoimmunity, including STIM1, which was further validated using co-immunoprecipitation. Our studies reveal novel proteins and molecular pathways that interact with Kv1.3 channels in adaptive (T-cell) and innate immune (microglia), providing a foundation for how Kv1.3 channels may regulate immune mechanisms in autoimmune and neurological diseases.

## INTRODUCTION

T-cells are major enactors of adaptive immunity (1). T-cells originate from thymocyte progenitors and mature into distinct subtypes, with varied molecular phenotypes, tissue localization and functions during normal development, response to infectious pathogens, autoimmunity and other diseases associated with chronic inflammation (2–4). Early T-cell precursor’s undergo sequential development to become double positive (DP) CD4+CD8+ precursor cells. T-cell receptor (TCR) signaling along with transcription factors regulate fate of double-positive thymocytes to adopt either CD4+ or CD8+ lineages (5, 6). Within CD4+ and CD8+ T-cell compartments, a wide range of naive, effector, memory and regulatory sub-classes exist (4). CD4+ T-cells differentiate into several subsets like Treg, Th1, Th2, Th17, Th9, Th22 etc., that can mediate helper functions with important roles in regulating B cell proliferation, antibody responses and functioning of CD8+ cells (7, 8), while CD8+ T-cells mediate cytotoxic functions (9). Naive CD4+ and CD8+ T-cells become activated short-lived effector T-cells upon antigen presentation, and a subset of these adopt longer-lived memory profiles, including central-memory (TCM) and effector-memory (TEM) phenotypes. These memory T-cells provide long-term immunity to infectious pathogens by responding with strong reactions on antigen re-encounter (9–11). Unique molecular characteristics and pathways that regulate functions of distinct T-cell subsets, provide avenues to T-cell sub-class-specific modulation, without impacting other classes (12).

T-cell activation involves a cascade of signaling events, initiated by interaction of the TCR with specific peptide ligands presented by antigen presenting cells (APC). Downstream signaling triggers Calcium (Ca^2+^) release from the endoplasmic reticulum (ER) to the cytosol (13). Sensing the depletion of ER Ca^2+^ levels, STIM1, an ER membrane protein, undergoes conformational change, it oligomerizes and expand towards the plasma membrane where it interacts with ORAI1 channels, leading to channel opening and Ca^2+^ influx, which then regulates T-cell function as well as differentiation (14). Pathways that regulate Ca^2+^ flux are therefore critical determinants of T-cell activation. Blocking Ca^2+^ leads to impairment of T-cell activation, proliferation, migration, cytokine production and effector functions (15). Potassium (K⁺) channels play a critical role in regulating this Ca²⁺ signaling by maintaining the electrochemical gradient and the negative membrane potential needed for sustained Ca²⁺ influx (16). Regulation of Ca^2+^ entry in T-cells is a major point of scientific investigation in the field of T-cell mediated human diseases, which span infectious illnesses, systemic autoimmunity (*e.g.* multiple sclerosis, rheumatoid arthritis), chronic diseases where altered T-cell functions have been identified (*e.g*. atherosclerosis or cancer), as well as neurodegenerative diseases (17). Many genetic risk factors associated with autoimmune diseases encode proteins that regulate immune signaling via Ca^2+^ flux pathways, *e.g.* STIM1, PTPN22, RUNX2 (18, 19) (https://www.ebi.ac.uk/gwas/efotraits/EFO_0005140).

While some Ca²⁺ influx pathways are shared across T-cell subtypes, the regulators of Ca²⁺ flux and signaling are unique to each subtype. For example, activated TEMs and Th17 cells predominantly express the voltage-dependent Kv1.3 potassium channel (Kv1.3) which maintains the potential gradient for Ca^2+^ flux, while TCM and Treg, utilize KCa3.1 channels to regulate their Ca^2+^ gradient (20–22). More than 100 human diseases are recognized as T cell-mediated autoimmune diseases, and among these, several are TEM-mediated, including type 1 diabetes mellitus, multiple sclerosis, and rheumatoid arthritis, being amongst the most prevalent (17). Effector memory T-cells from patients with TEM-mediated autoimmune diseases express high levels of Kv1.3 channels as compared to healthy individuals (20, 23). Notably, Kv1.3 channel blockade or deletion makes mice resistant to autoimmunity in disease models and promotes an immunoregulatory phenotype in T-cells (24). These studies highlight critical roles of Kv1.3 channels in regulating T-cell function and autoimmunity.

Kv1.3 channels, encoded by the *KCNA3* gene, were first described in T-cells in 1984 and have since been investigated as targets for immunosuppression (25, 26). The Kv1.3 channel is a voltage-activated K^+^ channel which exists as a homo-tetramer, with each monomer consisting of six transmembrane alpha helices, connected to each other through intra and extracellular loops (27). The channel opens following membrane depolarization (half-maximal activation potential typically around −30 mV), which can occur following Ca^2+^ entry in T-cells (28). K^+^ efflux via Kv1.3 channels restores membrane potential that facilitates sustained Ca^2+^ entry. Kv1.3 shows C-type inactivation and use-dependence with frequent depolarizations, both distinct characteristics of Kv1.3 channels as compared to other voltage-gated potassium (Kv) channels (29). The N and C-termini of Kv1.3 are cytosol-facing, and contain motifs that allow the channel to interact with cytosolic and membrane-associated proteins and signaling pathways (30, 31). These interactions link Kv1.3 channels to other immune pathways, raising the possibility that proteins interacting with or proximal to Kv1.3 channels can regulate channel function, and vice versa, where Kv1.3 function may regulate the function of interacting/proximal proteins and pathways. Cell surface receptors such as insulin-like growth factor (IGF) and epidermal growth factor (EGF) receptors, have indeed been shown to regulate Kv1.3 localization and activity via phosphorylation (32). IGF-1 increases K^+^ currents and up regulates Kv channel turnover in cells through PI3-kinase, PDK1 and SGK1 signaling cascades (33). EGFR activation recruits SRC family kinases or ERK1/2 signaling which modulates channel kinetics either by reducing activity or inducing channel endocytosis (34). Conversely, Kv1.3 channel function has been shown to impact interferon-mediated STAT1 activation in myeloid cells (35) as well as to interact with STAT3 in melanoma cells (36). Beyond regulation of K^+^ efflux and plasma membrane potential, Kv1.3 channels have been identified in the inner mitochondrial membrane, where they may regulate metabolism and apoptosis (37). Accordingly, apoptosis-regulatory proteins such as BAX have been identified to physically interact Kv1.3 channels in the mitochondrial inner membrane (38). Recently, Kv1.3 channel localization to the nuclear membrane in cancer cells has also been identified, suggesting that Kv1.3 may regulate membrane potential across several cellular compartments (39).

In T-cells, Kv1.3 channel functionally couples with β1-Integrins and shows molecular proximity with the CD3 complex (21, 40, 41). These co-localizations of Kv1.3 channels with signaling molecules suggest that Kv1.3 function may not be restricted to membrane potential regulation, but that the channel might also directly affect signaling proteins. Recently, proximity labeling approaches have been used to identify protein interactors of N and C-termini of Kv1.3 in mammalian cells, including HEK 293 embryonic kidney cells and BV-2 microglial cells (35, 36).., These studies showed that the N-terminus of Kv1.3 interacts with proteins involved in channel trafficking and localization whereas C-terminal-associated proteins regulate immune signaling, and several C-terminal interactions are dependent on the PDZ-binding domain of Kv1.3 channel. Although many studies have focused on the role of Kv1.3 channels in T-cell-mediated immune responses, not much is known about signaling pathways regulated by Kv1.3 channel in T-cells.

In this study, we aim to define the proximity-based protein interactome of Kv1.3 channels in T-cells. Our goal was to identify interactors that are unique to N and C-terminal domains of Kv1.3, and further identify proteins that are dependent on the PDZ-binding domain of C-terminus. Importantly, we also aimed to identify Kv1.3 interactors that are unique to T-cells as compared to microglia and nominate Kv1.3-associated pathways of importance to T cell-mediated autoimmune diseases. We employed a proximity labeling technique in which biotin ligase TurboID is fused to N or C-termini of Kv1.3 in human Jurkat T cells (JTC), so that proteins in proximity (10-30 nm) to Kv1.3 channels are biotinylated (42). As opposed to traditional co-immunoprecipitation methods that enrich highly stable protein-protein interactions, the TurboID approach allows labeling and quantification of both stable as well as transient but functionally important interactions. After enriching biotinylated proteins from whole cell lysates, we applied label-free quantitative mass spectrometry (LFQ-MS) of biotinylated proteins. We chose JTC, an immortalized line of human T lymphocytes, as our model system of choice for these studies because they endogenously express Kv1.3 channels and therefore contain cellular machinery for channel localization and function and recapitulate effector functions of activated T-cells (43). We used lentiviral transduction to generate T-cell lines stably expressing N and C-terminal TurboID fusions with Kv1.3, as well as a PDZ-binding domain-deleted C-terminal fusion. We used electrophysiology and flow cytometry to verify functional Kv1.3 channel presence in T-cells before LFQ-MS studies. This approach allowed us to define the proximity-based interactome of Kv1.3 channels in T-cells, including proteins that preferentially interact with N and C-termini respectively. The N-terminus of Kv1.3 interacts with proteins involved in mRNA processing and transcriptional regulation whereas the C-terminus interacts with proteins that regulate PDZ-dependent endocytic functions and maintains cell junction and synaptic organizations. We also identified Kv1.3 channel interactors that are unique to T-cells, as compared to microglia, as well as over 100 Kv1.3 interactors with causal roles in autoimmune diseases such as STIM1. Using co-immunoprecipitation, we validated the physical interaction between Kv1.3 and STIM1 in T-cells.

## MATERIALS AND METHODS

### Reagents

All key reagents used in this study are listed in **Table 1**.

**Table 1.** Reagents List.

| Reagent | Manufacturer | Catalog # |
| --- | --- | --- |
| RPMI-1640 | Gibco | 11875085 |
| Penicillin-Streptomycin | Gibco | 15140-122 |
| Fetal Bovine Serum (FBS) | Gibco | 26140-079 |
| Polybrene | Sigma-Aldrich | S2667 |
| Puromycin | Gibco | A11138-02 |
| Biotin | Sigma-Aldrich | B4639-100mg |
| 1x fixation buffer | eBioscience | 00-8222-49 |
| 1x permeabilization buffer | eBioscience | 00-8333-56 |
| Streptavidin, Alexa-Fluor 488 conjugate | Invitrogen | 21832 |
| Unstained OneComp ebeads compensation beads | eBioscience | 01-111-42 |
| HALT protease & phosphatase inhibitor cocktail | Thermofisher | 1861284 |
| Pierce streptavidin magnetic beads | Thermofisher | 83817 |
| BCA colorimetric assay | Thermofisher | 23227 |
| Bovine Serum Albumin Standards | Thermofisher | 23208 |
| 4x laemmli buffer | Bio-Rad | 1610747 |
| BOLT 4%-12% Bis-Tris Gels | Invitrogen | NW04125BOX |
| 250kDa Protein Ladder | BioLabs | P7719S |
| iBLOT mini stack system | Invitrogen | IB23002 |
| StartingBlockT20 | Thermofisher | 37543 |
| Streptavidin, Alexa-Fluor 680 conjugate | Invitrogen | S32358 |
| Silverstain | Thermofisher | 24612 |
| lysyl endopeptidase | Wako | 125-05061 |
| Trypsin | Promega | V5111 |
| HLB columns | Oasis | 186003908 |
| Reagent A | Thermofisher | 23222 |
| Reagent B | Thermofisher | 23224 |
| Milliplex® MAP Multi-Pathway<br>Magnetic Bead 9-Plex - Cell<br>Signaling Multiplex Assay | Millipore Sigma | 48-680MAG |
| NP-40 Lysis Buffer | Thermofisher | J60766.AP |
| Protein G Dynabeads | Thermofisher | 10004D |
| Anti-V5 tag antibody | Cell Signaling Technology | 80076S |
| Anti-V5 tag antibody | Abcam | ab206566 |
| Anti-STIM antibody | Cell Signaling Technology | 4916S |
| Anti-CD3 epsilon | Cell Signaling Technology | 85061S |
| Anti-NF-ATc1 antibody | BD Biosciences | 556602 |
| Anti-calnexin antibody | Abcam | ab22595 |
| Anti-Lamin B1 antibody | Cell Signaling Technology | 12586S |
| Anti-Tubulin antibody | Millipore | MAB1864 |
| Beta actin polyclonal antibody | Thermofisher | PA1-183-HRP |
| PAP-1 | Synthesized in Wulff laboratory at the<br>University of California, Davis. | N/A |
| ShK-223 | AmbioPharm | Api2651 |

### Plasmid and LV design

We utilized three previously validated plasmids that contained different human Kv1.3-TurboID fusion constructs, without additional modifications (35). The Emory Cloning core transformed the V5-TurboID-NES plasmid (AddGene, #107169) using a competent E. coli strain (DH5α) according to the manufacturer protocols. For purification of the plasmid DNA, QIAfilter Plasmid kits (Midi prep kit; Qiagen; catalog no.: 12243) were utilized following the manufacturer’s protocol. Restriction sites were introduced via the PCR primers and the V5-TurboID-NES sequence was subcloned into pCDH-EF1-MCS-BGH-PGK-GFP-T2A-Puro (CD550A-1) and sequenced (**Table 2**). A 15 amino acid linker that was GS rich (5’-GGCGGAGGGGGCTCA-3’x3) was inserted between *Kcna3* and a V5 tag followed by TurboID on either the 3’ or 5’ end of the *Kcna3* gene. A truncated form of Kv1.3 was also created, lacking a C-terminal TDV amino acid residues (Kv1.3 C-terminal⊗PDZ). These three residues are essential for Kv1.3 function and localization (44). All Kv1.3 fusion constructs included a V5-tag that was immediately upstream to the TurboID construct, to allow for TurboID-based proximity proteomics, in addition to V5-based co-immunoprecipitation studies. Together, this strategy allowed for identification of proteins that are proximal to Kv1.3 (within the labeling radius of fused TurboID) such as interactions that are indirect or transient, as well as direct and more stable physical interactions (via V5 co-IP). For all three constructs, puromycin resistance was used as a selection marker and DNA sequencing was used to confirm plasmid orientation and correct insertion.

**Table 2.**
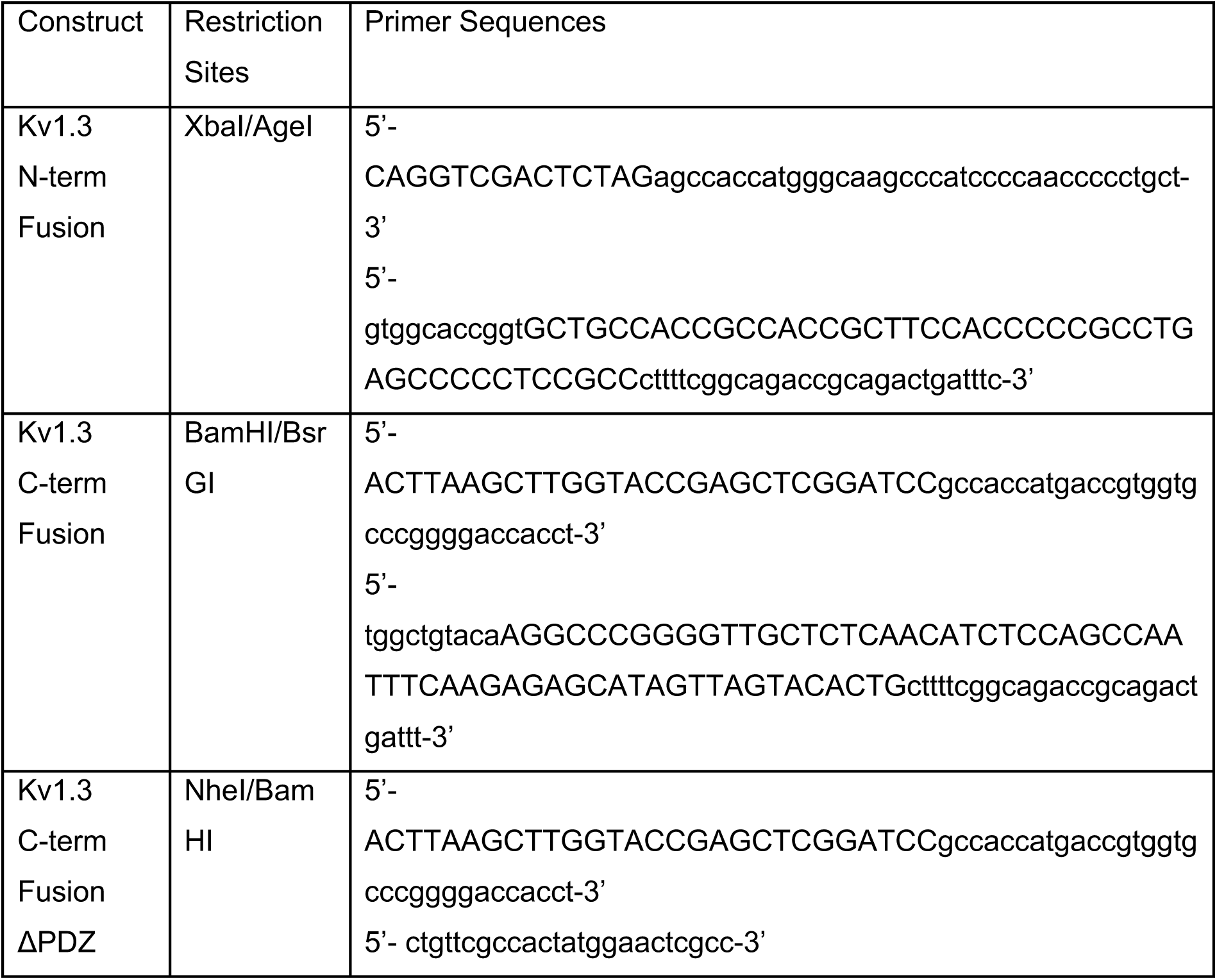
Constructs Designed for experiments conducted.

As previously described (35), plasmids were packaged into lentivirus (LV) by the Emory University Viral Vector Core and purified. The core utilized HEK-293FT (Invitrogen) cells for transfection. The cells were approximately 70-80% confluent in complete medium (4.5 g/L Glucose and L-Glutamine containing DMEM supplemented with 10% FBS and 1% Pen-Strep) and incubated at 37 ° C, 5% CO_2_ prior to transfection. The day of transfection, a 2 mL of a DNA and polyethyenimine (PEI) mixture was added dropwise to the HEK-293 cells then incubated for 48 h before harvesting.

Plasma purification was completed on supernatants, containing lentivirus 48 h and 72 h post-transfection following the protocol described in Bowen *et al* (35)Supernatants were centrifuged at 500 x g for 5 min at 4 ° C and passed through a 0.45 μm low protein binding filter. The 200 μL of supernatants were centrifuged at 28,000 rpm for 2 h at 40 ° C. The virus particles were resuspended in 500 μL of PBS and incubated on ice for 30 min. Resuspended virus particles were loaded into a 12 mL of SW 41 tube with a 3 mL of 20% sucrose cushion, and centrifuged at 28,000 rpm for 2 h at 4 ° C. The virus pellet was resuspended in 60 μL of PBS.

To transduce Jurkat T-cells (JTC), purified virus was added to cells at a multiplicity of infection (MOI) of 10 and 8 μg/mL polybrene for 24 h. Cells were collected in media and centrifuged at 125 x g for 5 min. Media containing LV was aspirated and cells were grown in RPMI-1640 media (10% FBS, 1% Penicillin/streptomycin) for 5 days. Puromycin was added at 2 μg/mL for 7 days.

### Cell Culture

JTC Clone E-6 cells were obtained from ATCC (Lot 70029114). Cells were maintained in RPMI-1640 (10% FBS and 1% Pen/Strep) at 1 million cells in 100 mm dish for protein lysis. Cells were allowed to stabilize for 24 h prior to experiment.

For channel blocking, cells were treated with PAP-1 (100 nM) or ShK-223 (200 nM) for 24 h. Supernatant was collected for cytokine analysis and cell pellets were lysed for other analysis. For the biotinylation of proteins, JTCs were exposed to 200 μM Biotin, which was added to the media for 1 h.

### Cell Lysis and protein processing

JTCs were collected in media and centrifuged at 125 x g for 5 min, then resuspended in 1 mL cold 1x PBS prior to centrifugation at 800 x g for 5 min at RT (n=3/experimental group). Cell lysis was completed exactly as previously published (35). Lysis was completed in 8 M urea in Tris-NaH_2_PO_4_ with HALT Protease inhibitor (1:100) and probe sonicated three times at 30% amplitude for 5 s each, with 10 s breaks between pulses. Lysates were centrifuged at 15000 x g for 15 min. Supernatants, containing solubilized proteins, were processed for western blot, streptavidin affinity purification (AP) and Mass Spectrometry (MS). Pellets using this method are minimal, resulting in most proteins being present in the eluate.

### Flow Cytometry

JTC cells were collected in growth media in a 15 ml centrifuge tube and centrifuged at 500 x g for 5 min. The supernatant was discarded, and cells were then further washed in the same tube twice by centrifuging the tubes using ice-cold 1x PBS at 800 x g for 3 min (n=3/group). The cells were then further transferred into the flow tube. Before fixation cells were processed on ice and by using cold PBS. To confirm biotinylation, cells were fixed with 1x fixation buffer (Thermo Fisher Scientific; Cat# 00822249) for 30 min, then washed three times with 1x PBS and centrifuged at 800 x g for 3 min. Cells were permeabilized for 30 min using 1x permeabilization buffer (Thermo Fisher Scientific; Cat# 00822249) and then incubated with Streptavidin-Alexa Fluor 488 conjugate (Thermo Fisher Scientific; Cat#S32354, 1:500 in permeabilization buffer) for 1 h in the dark. After incubation, the cells were washed in 1x PBS three times. After the last wash, 200 µL of PBS was added. Unstained OneComp beads and unstained cells were used as negative controls. Beads stained with Alexa fluoro-488 (Thermo Fisher Scientific; Cat# A-11001) were used as a positive control. Flow cytometry data was collected on the BD Aria II instrument and analyzed using the Flow Jo software (v10.10.0).

### Electrophysiology

JTCs were grown in RPMI 1640 medium (Sigma-Aldrich, St. Louis, MO) containing 10 % heat-inactivated FBS, 10 mM HEPES and 2 mM glutamate at 37 °C in a 5 % CO_2_ humidified incubator. Electrophysiological recordings were performed on transfected HEK-293 cells or transduced JTCs plated on poly-L-lysine-coated glass coverslips. All measurements were performed using the whole-cell configuration of the patch-clamp technique at room temperature with an EPC-10 HEKA amplifier. Transfected HEK-293 cells were visualized by epifluorescence microscopy of green fluorescent protein by eEGFP-C1 plasmid co-transfection (Addgene, 2487, discontinued) and these cells were used as wild-type Kv1.3 channels for comparison with Kv1.3-TurboID fusions in JTCs. The Ringer solution used contained (in mM) 160 NaCl, 4.5 KCl, 2 CaCl_2_, 1 MgCl_2_, 10 HEPES, pH 7.4, and adjusted osmolarity of 300 mOsm. Patch pipettes were made from soda lime glass (microhematocrit tubes, Kimble Chase) and had resistances of 2 to 3 MΩ when filled with internal KF solution and submerged in the bath solution. The pipettes were filled with an internal solution containing (in mM) 145 KF, 1 HEPES, 10 EGTA, 2 MgCl_2_, pH 7.2, and 300 mOsm. Series resistance (at least 80%) and whole-cell capacitance compensation were used as quality criteria for electrophysiology. Voltage-dependence of activation was recorded using a 200-ms voltage step from −80 to +60 mV in 10 mV increments with an inter-pulse duration of 30 s, and analyzed as previously described (35). Use-dependency was determined using single pulse to +40 mV at a pulse frequency of 1 Hz for 10 pulses. The fractional current of the last pulse was normalized to the first pulse to determine the extent of cumulative (use-dependent) inactivation. Whole-cell patch-clamp data are presented as mean ± SD, and statistical significance was determined using a paired Student’s t test for direct comparison between wild-type JTC (without transduced Kv1.3) and JTC transduced with Kv1.3-TurboID fusion lentiviruses.

### Western Blot

Protein quantification and western blots were performed exactly as previously described (35). Briefly, proteins were quantified using a bicinchoninic acid assay (BCA) colorimetric assay (n=3). To evaluate biotinylation of proteins, 10 µg of protein was resolved on a BOLT 4%-12% Bis-Tris Gel. Proteins were transferred to a nitrocellulose membrane using a semi-dry transfer system. Ponceau staining for 2 min was used to determine loading efficiency. Blots were washed with 1x TBS, blocked for 30 min with StartingBlock and probed with streptavidin-Alexa fluoro-680 (1:10,000 in Blocking Buffer) for 1 h at room temperature protected from light. The membrane was then washed with 1x TBS and imaged using the Odyssey Li-COR system or Bio-Rad ChemiDoc MP Imaging system (Universal Hood III). Densitometric analyses were done using ImageJ software. Statistical significance was measured using one-way ANOVA followed by Tukey’s test in Prism 9 software.

### Luminex

Multiplexed phospho-protein quantification was conducted for 9 phosphorylated proteins involved in various signaling pathways using the Milliplex® MAP Multi-Pathway Magnetic Bead 9-Plex - Cell Signaling Multiplex Assay (CREB pS133, ERK pT185/pY187, NFκB pS536, JNK pT183/pY185, p38 Thr180/Tyr182, p70S6K Thr412, STAT3 pS727, STAT5A/B pY694/699, Akt pS473). Lysates were thawed on ice and centrifuged at 4°C for 10 minutes at 15,500 × g. Protein concentrations were normalized to 1.5 µg sample per 12.5 µL using Milliplex® MAP Assay Buffer for analysis. These protein concentrations were selected to ensure they fell within the linear range of bead fluorescent intensity versus protein concentration for detectable analytes.

For multiplex assay, background measurements were collected using assay buffer in the absence of biological samples. The average background values were subtracted from each sample measurement, and any negative values were set to zero. Heatmap for visualization were generated using z-scored data and the R package heatmap3. Statistical significance was assessed using one-way ANOVA followed by Tukey’s test, performed in Prism 9 software.

### Co-immunoprecipitation (co-IP) assay

Untransduced JTCs and N-terminal Kv1.3-TurboID cell lysates were prepared in NP40 cell lysis buffer followed by protein content estimation of cleared cell lysates by BCA assay. Equal amounts of each cell line’s extracts were incubated with 1 μg of V5 antibody overnight at 4°C. Protein G Dynabeads were added to each sample (10%, v/v) and incubated further for 1 h at 4°C. After three washes with NP40 lysis buffer, samples were recovered in 50 μl of 4x SDS gel-loading buffer and subjected to western blot as described above. Immunoprecipitated proteins and input samples were run simultaneously and probed for the presence of STIM1, CD3E, V5 tagged protein and actin using different antibodies. Co-IP of Kv1.3 with STIM1 and CD3E was checked in N-terminal Kv1.3-TurboID JTCs by using untransduced JTCs as control.

### Affinity purification of biotinylated proteins

Streptavidin affinity purification was performed on cell lysates, to capture biotinylated proteins. Affinity purification was performed in the same manner as Bowen *et al* (*35*) where, Pierce streptavidin magnetic beads were washed with RIPA buffer. 1 mg of protein lysate was suspended in RIPA buffer and incubated with streptavidin magnetic beads. Beads were washed to assure reduction of non-specific binding following the protocol from Bowen *et al* (*35*). The beads, with biotinylated proteins attached, were suspended in 80 μL of PSB. 8 μL of beads (10% of total) were resuspended in Laemmli buffer supplemented with biotin and DTT and prepared for Western blot and silver stain. Western blot was used to evaluate the presence of biotinylated proteins post-AP. Silver stain was utilized to evaluate total protein abundance, and thus efficacy of affinity purification of post-AP samples. Following western blot and silver stain, remaining protein bound beads were stored at −20 °C until on-bead digestion.

### Protein Digestion and Peptide Clean Up

Preparation of samples for mass spectrometry followed protocols outlined in Bowen *et al.* (35). For biotin enriched samples from the AP, samples were washed with 1x PBS and resuspended in 50 mM ammonium bicarbonate buffer (ABC). Samples were reduced utilizing dithiothreitol (DTT) and cysteines were alkylated by iodoacetamide (IAA). Proteins were digested with 0.5 µg of lysyl endopeptidase and 1 µg trypsin overnight. Samples were acidified and desalted using an HLB column. Samples were dried using cold vacuum centrifugation (SpeedVac Vacuum Concentrator).

For cell lysates, 100 μg of pooled cell lysates were utilized. Reduction and alkylation were the same as described above. Samples were diluted in ABC and digested with 2 µg of lysyl endopeptidase and 4 µg trypsin. Samples were acidified, desalted using an HLB column, and dried down using cold vacuum centrifugation (SpeedVac Vacuum Concentrator). Digestion protocol is also outlined in Sunna, *et al*. and Rayaprolu *et al*. (45, 46).

### Mass Spectrometry (MS)

Derived peptides were suspended in 0.1% trifluoroacetic acid and separated on a Water’s Charged Surface Hybrid column. Jurkat T-cells cell samples were run on an EVOSEP liquid chromatography system using the 30 samples per day preset gradient and were monitored on a Q-Exactive Plus Hybrid Quadrupole-Orbitrap Mass Spectrometer (Thermo Fisher Scientific). MS protocols used here have been outlined in prior publications and were used without modifications (Sunna *et al.* and Bowen *et al*. (35, 46). The mass spectrometry proteomics data have been deposited to the ProteomeXchange Consortium via the PRIDE (47) partner repository with the dataset identifier PXD059644.

### Protein Identification and Quantification

MS raw files were uploaded into the MaxQuant software (version 2.4.9.0), and JTC data were searched against the human Uniprot 2022 database (83,401 proteins) with the addition of the target sequence for TurboID. MaxQuant parameters used were exactly as previously published (35). Modifications were limited to a maximum number of five and include variable modifications: Methionine oxidation, protein N-terminal acetylation, and fixed modification: carbamidomethyl. The Maximum number of missed cleavages was set to 1. For peptide size, the maximum peptide mass was set to 6000 Da and minimum peptide length of 6 amino acids. Label Free Quantification (LFQ) minimum ratio count was set to one. Unique and razor peptides were used for peptide quantification. The minimum ratio count for protein quantification was set to 1. Re-quantification ensured the peptide matched with the identified protein. Identifications were matched between runs. To determine m/z ratio, Fourier transformed MS (FTMS) match tolerance was set to 0.05 Da. Ion Trap MS was set to 0.6 Da to evaluate gaseous ions. The false discovery rate (FDR) was set to 1%.

For both the biotin enriched samples and cell lysates, MaxQuant intensities were uploaded into Perseus (version 1.6.15). To obtain proteins that best match the peptides, potential contaminants, reverse peptides, and samples only identified by one site were all removed. Both datasets were log2 transformed. Data was imputed based on normal distribution for 50% missingness within an experimental group. Cell lysate data was matched to uniport gene names. To ensure that any differences observed are not due to unequal TurboID expression or unequal biotinylation, all LFQ-MS values from the AP dataset from Kv1.3-TurboID lines were normalized to TurboID protein abundance, while bulk proteome was normalized using quantile normalization (**Supp. Datasheet 1**).

### Protein-protein interaction (PPI) network analysis

The search tool for retrieval of interacting genes (STRING) (https://string-db.org) database, which integrates both known and predicted PPIs, can be applied to predict functional interactions of proteins (48). To seek potential interactions between DEPs, and for cellular compartmentalization of proteins, the STRING tool was employed. Active interaction sources, including text mining, experiments, databases, and co-expression as well as species limited to “Homo sapiens” and an interaction score > 0.4 were applied to construct the PPI networks. Cytoscape software version 3.6.1 was used to visualize the PPI network.

### Functional and pathway enrichment analysis of DEPs

Gene Ontology (GO) pathway enrichment analyses were conducted using Enrichr (http://amp.pharm.mssm.edu/Enrichr/) and ClusterProfiler R packages. The significant terms and pathways were selected with the threshold of adjusted *p*-value < 0.05.

### Data availability

All raw MS data generated in this study are available via ProteomeXchange with identifier PXD059644. Kv1.3 interactors in the BV-2 dataset was from an earlier published study by Bowen *et al.*, available at ProteomeXchange Consortium with the dataset identifier PXD049433 (35).

Membrane proteins data taken from EBI-GO_Membrane and Membranome databases (https://membranome.org/proteins and https://www.ebi.ac.uk/QuickGO/term/GO:0016020).

List of autoimmune disease associated risk genes was collected from EMBL-EBI databases (https://www.ebi.ac.uk/gwas/efotraits/EFO_0005140).

## RESULTS

### Validation of stably transduced Jurkat T-cells that express N- and C-terminal fusion of TurboID to Kv1.3 channels

JTCs are immortalized human T-cells, known to express Kv1.3 channels and Kv1.3-associated immune proteins, making this a valid *in vitro* model system for Kv1.3 channel proximity-based proteomics (43). Using Kv1.3-TurboID fusion, we generated JTC lines that stably expressed these fusion proteins. JTCs were transduced for 24 h with lentivirus (LV) containing one of three constructs: The human *KCNA3* gene fused to TurboID on the N-terminus, Kv1.3 fusion with TurboID on the C-terminus, and a Kv1.3-TurboID C-terminal fusion with the PDZ-binding domain removed (**Fig. 1A**). Untransduced JTCs were utilized as a negative control. Following puromycin selection for seven days, transduced cell lines were exposed to biotin for 1 h to biotinylate proteins within proximity to Kv1.3 (**Fig. 1B**). Flow cytometry and western blot show JTCs transduced with Kv1.3-TurboID contain biotinylated proteins at much higher levels than compared to the endogenously biotinylated proteins in control JTCs (**Fig. 1 C and D**). The loading of proteins on western blot were evaluated via ponceau staining (**Supp Fig. 1A**). We then enriched biotinylated proteins using streptavidin beads. Biotinylated proteins and endogenously biotinylated proteins were primarily enriched from JTC-TurboID cells as compared to control JTCs, evidenced by silver stain detection of all enriched proteins (**Supp Fig. 1B**) and WB to detect biotinylation (via streptavidin conjugated-fluorophore) (**Supp Fig. 1 C**). These biochemical results confirm that TurboID is present in the JTC transduced with Kv1.3-TurboID. Furthermore, the unique banding patterns indicate that there are likely different proteins being affinity purified for each construct.

**Figure 1:**
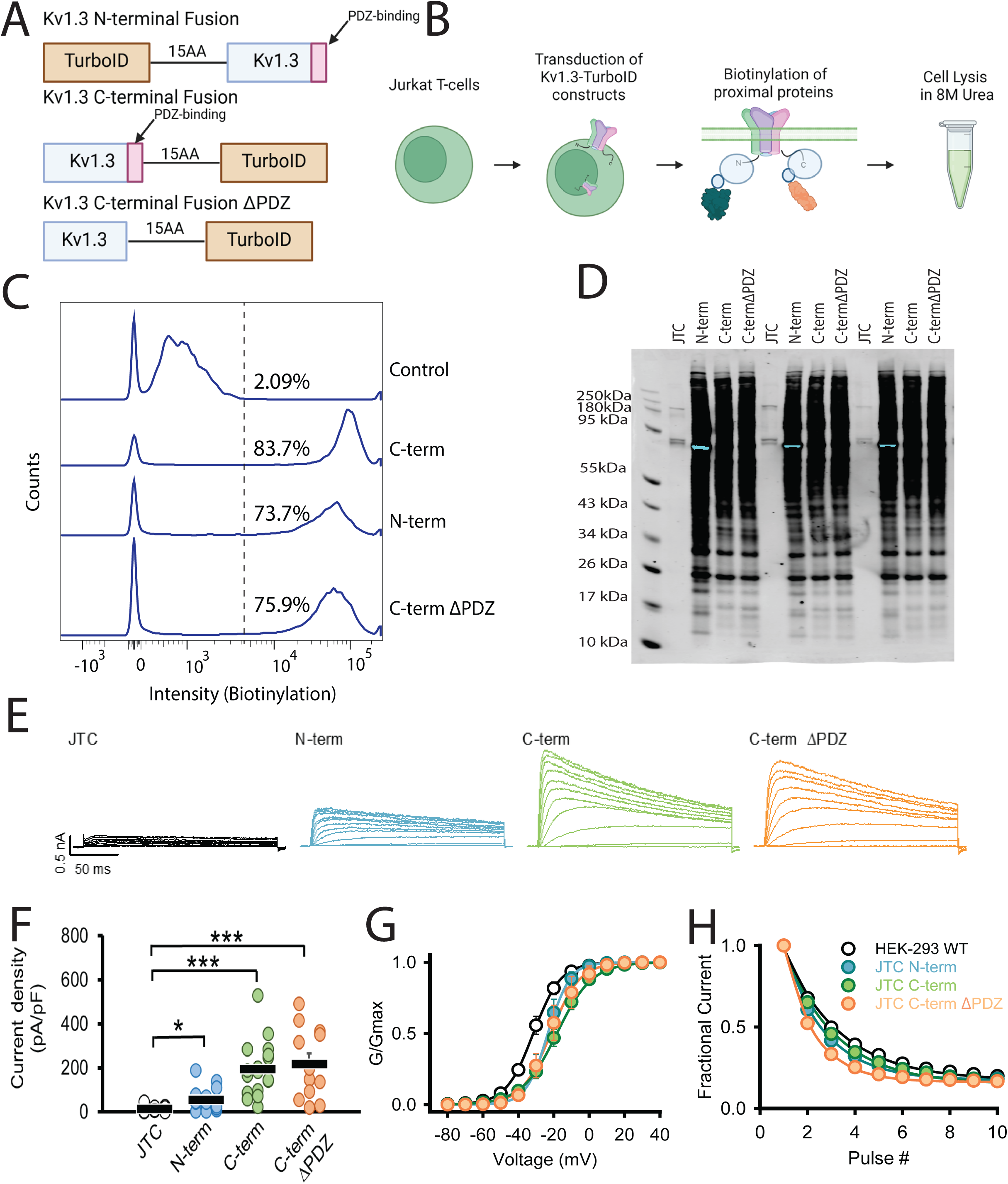
TurboID biotinylates proteins in Kv1.3-TurboID transduced Jurkat T-cells. **(A)** Constructs utilized for transduction into Jurkat T-cells (JTC). Three constructs were used: a fusion of TurboID to the N-terminus of Kv1.3, a fusion of TurboID to the C-terminus of Kv1.3, and the C-terminal fusion with the PDZ-binding domain deleted. **(B)** Schematic of experimental design. Jurkat T-cells were transfected with Kv1.3-TurboID fusion constructs then exposed to biotin. Cells were then lysed in 8M urea. **(C)** Flow cytometry analysis shows transduced cells have higher biotin present than control cells (n=3). **(D)** Western blot utilizing streptavidin-680 shows distinct biotinylation patterns in each transduced cell line (n=3). **(E)** Representative currents recorded from wild-type JTC activated with concanavalin A (5 µg/mL), JTC transduced with N-terminal fusion Kv1.3 (N-term), Kv1.3 C-terminal fusion (C-term), and C-terminal fusion with the PDZ-binding domain deleted (C-term⊗PDZ) constructs. Currents elicited by a voltage step protocol from holding potential of −80 mV to +60 mV in 10 mV increments. **(F)** Current density calculated from peak currents induced by the +40-mV depolarization step in wild-type JTC (n=18), N-term (n=17), C-term (n=18), and C-term⊗PDZ (n=12). **(G)** Conductance-voltage relationship depicting voltage-dependence of activation of WT Kv1.3 construct transfected in HEK-293 (V_1/2_ = −31.4 ± 1.7 mV, n=8), and JTC transduced with N-term (V_1/2_ = −23.6 ± 6.0 mV, n=6), C-term (V_1/2_ = −18.8 ± 6.0 mV, n=6), and C-term⊗PDZ (V_1/2_ = −21.1 ± 10.2 mV, n=7) fusion constructs. **(H)** Fractional currents showing no changes in use-dependent current reduction of Kv1.3 currents in JTC transduced with N-term (n=9), C-term (n=10), and C-term⊗PDZ (n=8) in comparison with wild-type Kv1.3 transfected in HEK-293 (n=8). Statistical significance denotes p < 0.05 (∗) and p < 0.001 (∗∗∗). BioRender software was used to generate Fig 1B.

We next utilized whole-cell patch-clamp electrophysiology to evaluate the functional expression of recombinant Kv1.3 channels at the cell surface of JTCs. All three transduced Kv1.3-TurboID JTC lines (N-terminal, C-terminal and C-terminal ⊗PDZ) exhibited robust current expressions with comparable amplitudes and densities (**Fig 1E**). Analysis of voltage-dependent activation and use-dependent inactivation revealed similar properties across all constructs (**Fig 1E and F)**. However, all displayed a slight shift in voltage-dependence of activation toward a more positive potential and a modest decrease in residual fractional current compared to WT Kv1.3 transiently expressed in HEK-293 cells (**Fig. 1 G and H**).

To further assess whether Kv1.3 over-expression impacts key immune signaling pathways in JTCs, we checked levels of NFAT activation, in nuclear and cytosolic fractions of JTCs, using the N-terminal Kv1.3-TurboID JTC line. We also treated control and transduced JTCs with the Kv1.3 inhibiting small molecule PAP-1 (100 nM) and the peptide ShK-223 (200 nM) for 24 hours to block channel activity. Western blot of nuclear and cytosolic fractions of treated JTCs showed that NFATc1 was primarily present in the nuclear fraction of cells, with no differences between treated control JTCs and N-terminal Kv1.3-TurboID JTCs, and no observed effect of PAP-1 and ShK-223 **(Supp Fig. 2A and B)**. Therefore, we concluded that neither Kv1.3 channel transduction nor Kv1.3 blockade affected NFAT translocation to the nucleus in JTCs. We also found TurboID localization by probing cell fractions with V5-tag, which is attached to TurboID-Kv1.3 constructs, was primarily located to the cytosolic fraction **(Supp Fig. 2A and B),** with no observed effects of Kv1.3 blockade. The purity of nuclear and cytosolic fractions was confirmed by probing the blot with Lamin B1, a nuclear-specific protein, and Calnexin, an ER-specific protein, along with measuring β-actin and Tubulin levels, which were enriched in cytosolic fractions. Similarly, we evaluated the impact of channel transduction and blockade on major cell signaling proteins by multi-plexed Luminex assay. There was no change with channel-transduced activation of pCREB, pAKT and pp38 signaling, while levels of pJNK, pERK, pNF-kB, pSTAT3 and pSTAT5 were increased in cell lysates **(Supp Fig. 2C and Supp. Datasheet 2),** but there was no effect of channel blockade on it. No impact of channel blockers signifies that the increase in the cytokines and signaling proteins is not an outcome of Kv1.3 channel transduction. As in Luminex assay, streptavidin fluorophore binds with biotinylated analyte, it is possible that the much higher number of biotinylated proteins in N-terminal Kv1.3-TurboID JTC line interfere with Luminex assay, resulting in apparent increase in these proteins. Overall, these biochemical and electrophysiological data confirm functional Kv1.3 channel expression in our Kv1.3 JTC lines, without effects of Kv1.3 over-expression and/or cytosolic proteomic biotinylation on T cell activation, as reflected by unchanged NFAT activation levels. Importantly, channel properties were not significantly impacted by TurboID fusion, and the Kv1.3 channel densities observed by over-expression were similar to Kv1.3 channel densities observed in TEM cells *in vivo* in autoimmune diseases (23).

### Characterization of the Kv1.3 channel interactome in JTCs using TurboID proximity labeling

After validating our JTCs lines as described above, we proceeded to identify Kv1.3 channel interacting proteins in JTCs using LFQ-MS. We evaluated the streptavidin affinity-purified (AP) proteome and compared all Kv1.3-TurboID JTCs to the non-TurboID control JTC group. This initial approach was used to provide a broad survey of proteins in proximity to the Kv1.3 channel complex, regardless of preferential interactions with the N-terminal or C-terminal. Given the labeling radius of TurboID and the tetrameric structure of Kv1.3 channels, we predicted that the interactors using our approach would be identified across both N-terminal and C-terminal proximity proteomes while a smaller proportion will be specific to each terminus. Based on the level of enrichment comparing TurboID to non-TurboID groups, and imposing statistical thresholds, top-interacting proteins were nominated. In addition to analyzing the AP proteome, we also assessed the bulk proteome of each group, to ensure that any differences at the level of AP were not due to whole-cell-level proteomic changes due to LV transduction and/or Kv1.3 channel over-expression.

In the AP dataset, LFQ-MS across all samples yielded 2,571 proteins, with higher total protein intensity and number of identified proteins from TurboID groups as compared to non-TurboID control JTCs. Principal component analysis (PCA) of the AP proteomes showed group-wise separations (**Control, N-Terminal**, **C-Terminal,** and **C-terminal⊗PDZ**). The first principal component (PC1), accounting for 76% of the total variance, separated the experimental groups from the non-TurboID controls, confirming that the AP proteome from TurboID JTCs is systematically distinct from non-TurboID control JTCs. The second principal component (PC2), explaining 10% of the variance, highlighted distinctions between the N-terminal and C-terminal groups, while separation between C-terminal and C-terminal⊗PDZ groups was not observed based on PC1 and PC2. This demonstrates that while PC1 captured the primary variability related to condition, PC2 revealed secondary variations, possibly linked to specific protein modifications or subgroup-specific effects (**Fig. 2A**). To identify Kv1.3 channel interacting proteins, we conducted differential enrichment analysis (DEA) between controls and Kv1.3 labeled groups in both the AP and bulk proteome datasets. In the AP proteome, we identified 1,947 differentially expressed proteins (DEPs) that were biotinylated and therefore enriched in the Kv1.3-TurboID group as compared to non-TurboID group. Of these, 1,845 DEPs showed >4-fold enrichment with p-values <0.05 (Supp Table 1A), including CLINT1, PICALM, DOK1, GEN1 as well as Kv1.3 and TurboID, as would be expected based on our experimental design (**Fig 2B**). We applied the same threshold for all further DEA. 918 DEPs were identified at the level of bulk proteomes in Kv1.3-TurboID JTCs, including 453 decreased (*e.g.* FSCN1, PCCA, EIF5A) and 465 increased (*e.g.* COG1, CNN2, YKT6, ACBD3) proteins (**Fig 2C**). This suggests that LV transduction and/or Kv1.3 over-expression does modify the proteome of JTCs in our *in vitro* system.

**Figure 2:**
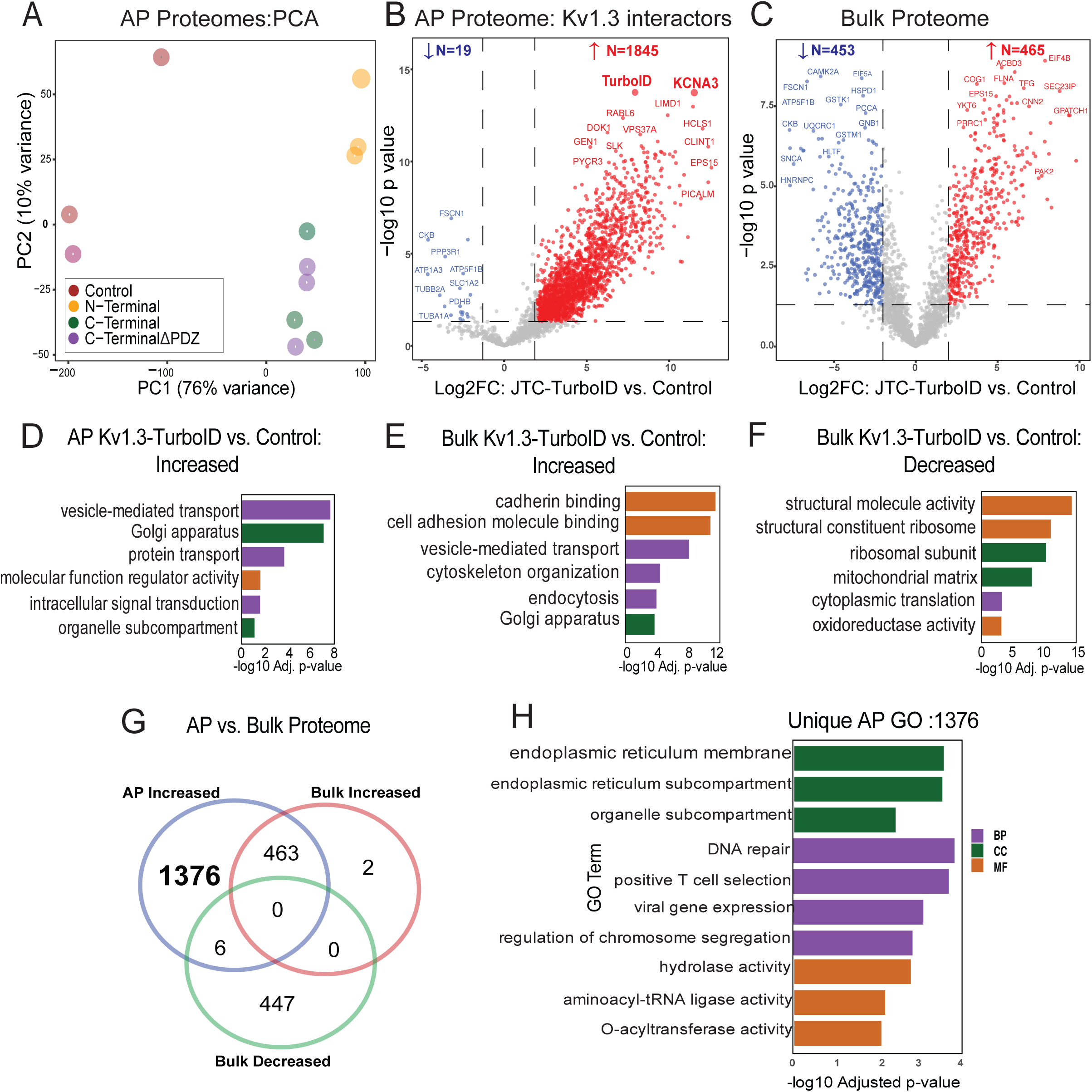
Comparative analysis of biotinylated proteins in JTC Bulk vs. JTC affinity purified samples. (**A)** Principal Component Analysis (PCA) of proteins from Kv1.3-TurboID transduced Jurkat T-cells and control samples (n=3). PC1 and PC2 explain 76% and 10% of the variance, respectively. Distinct clustering of sample groups reflects significant differences in each sample proteome. **(B)** Volcano Plot for JTC Affinity Purified (AP) Samples: Differential Enrichment Analysis (DEA) between all Kv1.3-TurboID fusion and control in affinity purified (AP) samples, showing 1845 proteins enriched with Kv1.3 channel transduction. **(C)** Volcano Plot for JTC Bulk Samples: DEA of all Kv1.3-TurboID fusion and control in bulk samples showing similar number of proteins increased (465) and decreased (453) with Kv1.3 channel transduction at bulk level. **(D)** Gene Ontology (GO) enrichment for increased proteins in AP samples identifies Kv1.3 enriched proteome involved in protein transport and signal transduction pathways. **(E)** GO Enrichment for increased proteins in JTC Bulk shows cadherin binding and vesicle mediated transport pathways enrichment with Kv1.3-TurboID fusion in bulk sample. **(F)** GO Enrichment analysis of decreased proteins in JTC bulk samples DEA shows downregulation of RNA processing, cytosolic translation, and mitochondrial function with Kv1.3 channel in bulk samples. **(G)** Venn diagram illustrates the overlap and unique proteins identified across JTC bulk and JTC affinity purified samples, with 1376 Kv1.3 enriched proteins unique to AP proteome, and overlap of 469 proteins with DEPs in bulk samples. **(G)** GO analysis for unique AP-Kv1.3 interacting proteins identified in Fig 2G emphasizes enrichment of processes associated with endoplasmic reticulum organization, membrane-associated trafficking, and Golgi-mediated vesicular transport, suggesting role of Kv1.3 in regulating protein localization and intracellular membrane dynamics. Complete protein list for each analysis is given in Supp. Datasheet 3.

Gene Ontology (GO) analysis of DEPs in the AP dataset (n=1845) showed enrichment in vesicle-mediated transport protein (FLNA, CORO1A, RAB6A), intracellular transport (GSK3A, RAB7A, COPB1), regulator activity (TAB1, ATP6AP1, MARK2) and Golgi-localized proteins (TMED10, FASN, CLCN3) (**Fig. 2D**). The presence of a high number of proteins associated with transportation, along with Golgi and ER proteins, suggests potential interactions during Kv1.3 protein processing and trafficking to the cell surface. In contrast to the AP proteomes, the whole-cell proteomes of TurboID-Kv1.3 JTCs showed increased levels of proteins involved in cadherin binding (PTPN11, GOLGA3, DLG1), cell adhesion (SCRIB, CTTN, ATP2C1), vesicle membrane (RAB14, COPZ1, STX5) and vesicle-mediated transport (GRIPAP1, GAPVD1, FNBP1L) functions **(Fig. 2E)** while ribosomal (BYSL, WDR36), structural molecule activity (NUP214, TLN1, SPTBN1) and mitochondrial matrix proteins (SHC1, CCAR2) were decreased (**Fig. 2F**).

Given the observed proteomic differences in whole-cell proteomes of Kv1.3-TurboID and non-TurboID JTCs, we identified Kv1.3 interactors from the AP dataset that were not identified as DEPs at the whole-cell proteome level. This yielded 1,376 Kv1.3 interacting proteins in the AP dataset (**Fig 2G**). GO terms enriched in this list of 1,376 Kv1.3 interactors (**Fig 2H**) resulted in mostly similar observations from **Fig 2D**, along with distinct pathways such as positive T cell selection, and hydrolase activity, showing that the majority of Kv1.3-interacting proteins, and their associated ontologies, cannot be explained by baseline differences in protein levels purely due to Kv1.3 over-expression or LV transduction effects.

### Unique N-terminal and C-terminal interactomes of Kv1.3 channels in JTCs

Past studies have suggested that the N-terminus and C-terminus of Kv1.3 channels interact with distinct protein partners, suggesting that these channel domains functionally interact with distinct molecular complexes and immune signaling pathways within immune cells (35, 36). To identify Kv1.3 channel interacting proteins that preferentially interact with the N-terminal or C-terminal of Kv1.3, we first compared N-terminus or C-terminus AP proteomes with non-TurboID controls to identify proteins unique to either N-terminal or C-terminal AP interactomes. Next, we directly compared the N-term and C-term AP proteomes to identify DEPs between these interactomes. As compared to controls, there were 1,946 DEPs in proximity of the Kv1.3 channel’s N-terminus (**Fig. 3A**). GO analysis identified enriched proteins involved in processes such as vesicle-mediated transport (MYO1G, SNAP23, STXBP3), Golgi vesicle transport (AP3D1, SEC16A, SNX3), protein transport (IPO5, KPNA4, TNPO2) and microtubule organizing center (DDX3X, RGS14, FLOT1) (**Supp Fig. 3A**). Interestingly, many of the interacting proteins are components of centrosome and microtubule organizing centers. In contrast, the C-terminal Kv1.3 proteome contained 1,304 DEPs (**Fig. 3B**), including proteins involved in vesicle-mediated (CLTC, NPC1, GAK) and endosomal transport (VPS26A, ARL1, VCP) and nucleoside-triphosphate (DENND11, RGS12, IPO7) and GTPase regulator (EIF5, TSC2, RANBP1) activity. Enriched proteins were localized to the Golgi apparatus (PIK3C2A, SCAMP3, AP1G1) endosome (NPC1, ARPC2, STX7) and endosomal membrane (PLIN3, TOM1, EPS15) (**Supp Fig. 3B**). Consistent with our prediction, this analysis shows that majority of interactors of the N-terminus and C-terminus identified by the TurboID-fusion method, are overlapping. Yet, we found 672 proteins unique to the N-terminal proteome and 30 proteins unique to the C-terminal proteome (**Fig. 3C**). DEA of N-terminus vs C-terminus AP proteomes identified 431 DEPs, with 390 enriched in the N-terminus and 41 enriched in the C-terminus (**Fig 3D**).

**Figure 3:**
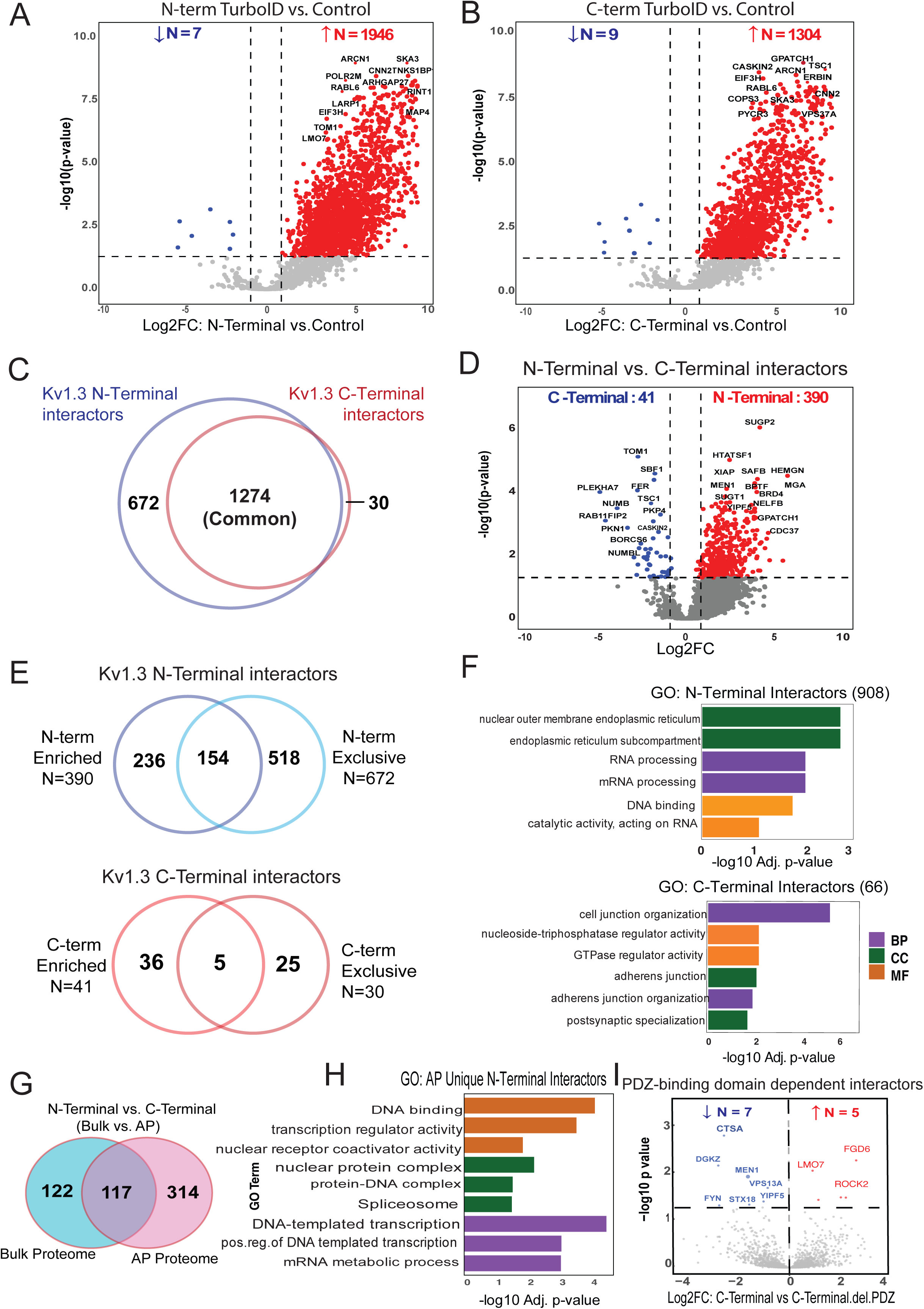
Distinct protein dynamics and enriched pathways in N-Terminal and C-Terminal interactors of Kv1.3 channel. **(A)** DEA of N-Terminal-TurboID and Control JTC, illustrates 1946 interactors of Kv1.3 channel’s N-terminal. **(B)** DEA of C-Terminal-TurboID and Control JTC, highlights 130 proteins enriched at Kv1.3 channel’s C-terminal. **(C)** Intersection of N-terminal (1946) and C-terminal (1304) interactors reveal 672 unique proteins associated with N-terminal and 30 with C-terminal. **(D)** DEA of N-terminal-TurboID and C-terminal-TurboID JTCs shows distinct N- (390) and C-terminal (41) interactors of Kv1.3 channel. **(E)** Venn diagram analysis intersecting N-terminal or C-terminal specific proteins identified in Fig 3C and D, shows 908 and 66 unique proteins for N-terminal and C-terminal, respectively. **(F)** GO term enrichment analysis for unique N- and C-terminal interactors identified in Fig 3E identifies protein processing function of N-terminal and cell signaling function of C-terminal. **(G)** Intersection of DEPs from N-terminal and C-terminal DEA at AP and bulk level shows 314 proteins differentially expressing only in AP proteome. Further analysis showed 310 and 4 are interactors of N-terminal and C-terminal, respectively. **(H)** GO term enrichment analysis for only AP proteome N-terminal interactors shows terminal role in transcription. **(I)** DEA between C-terminal and C-terminal with PDZ-binding domain removed shows 5 proteins increased and 7 decreased with domain deletion. All analyses are given in detail in Supp. Datasheet 4.

To identify differences in the N-terminal and C-terminal interactomes of Kv1.3, we overlapped unique N-terminal (672) and N-terminal enriched (390) interactors, to find total 908 proteins corresponding to N-terminal (**Fig 3E**). These N-terminal interactors included RNA/mRNA-processing proteins (RNMT, PRPF3, PLRG1, SART1, HTATSF1, NPM1) and ER membrane proteins (STING1, CDS2, TMX1). (**Fig 3F**). Similarly, for C-terminal we found 66 unique interactors (**Fig 3E**) which included proteins, mostly localized at cell adherens junctions (NF2, ADD1, PKP4), postsynaptic specializations (TSC1, ERBIN, LZTS1), and cell-cell junctions (FER, SIRT2, PKP4). Functionally, these proteins are involved in maintaining cell junctions (PLEKHA7, DOCK10, PEAK1, NUMB, NUMBL) and adherens junction organization (FER, ADD1) (**Fig 3F**).

To ensure that any N-terminus and C-terminus interacting proteins of Kv1.3 were not due to baseline effects of Kv1.3 over-expression or LV transduction, we compared DEPs of N-terminus vs. C-terminus in the AP proteomes (N=431) with those at the bulk (input) level (N=239). We found that the majority of DEPs identified as N-terminal or C-terminal interactors from the AP proteome (n=314) were not identified as DEPs at the whole-cell levels (**Fig 3G)**. Further GO analysis showed that 310 of these were interactors of the N-terminus, with mostly functions overlapping from **Fig. 3E**, and few unique such as DNA binding (BCL2, IMPDH2, TMPO) and DNA-templated transcription (ACTN4, TACC1, KAT7) (**Fig 3H**). Only four interactors were associated with the C-terminus, including Rho GTPase activating protein activator ARHGAP32 (49), PEAK1 (a kinase associated with cancer progression) (50), and proteins SNX17 and MTMR1, which show high affinity for the phosphatidylinositol 3-phosphate signaling molecule (51, 52).

In summary, our analyses show that the N-terminus of Kv1.3 channel interacts with proteins involved in RNA/mRNA processing and DNA transcription, ER processing and channel trafficking, while the C-terminus associates with proteins corresponding to cell junctions, synaptic organization and signaling.

### Proximity labeling nominates Kv1.3 interactors influenced by the PDZ-binding domain of the C-terminus

The PDZ-binding domain located at the C-terminus of the Kv1.3 channel plays a crucial role in its functional regulation, facilitating interactions with proteins like PSD95 and SAP97 (53, 54). This prompted us to investigate how deleting the PDZ-binding domain would affect the C-terminal interactome in JTCs. Upon comparing interactors of the full C-terminus with those of the C-terminalΔPDZ, we observed that deletion of the PDZ-binding domain reduced interactions with five proteins: FGD6, LMO7, ROCK2, TBC1D10B, and TUBB6—all of which are associated with cellular GTPase activity or pathways involving GTPases (55–59) **(Fig 3I)**. Interestingly, the PDZ-binding domain deletion also enhanced interactions with seven new proteins, including kinases FYN and DGKZ, lysosomal protease Cathepsin A (CTSA), and other proteins: YIPF5, VPS13A, STX18, and MEN1. Most of these new interactors are enriched in the ER, suggesting a shift in Kv1.3’s interaction profile toward proteins involved in cellular trafficking and signaling. In summary, deletion of the PDZ-binding domain from C-terminus of Kv1.3 affects major signaling interactions of the channel, along with introducing new interactions. PDZ-binding domain loss will thus potentially impact channel function.

### Comparative analysis reveals cell type-specific functions of Kv1.3 channel interactors in JTCs and BV-2 cell line

While Kv1.3 channels primarily regulate K^+^ transport, membrane potential, and Ca^2+^ flux, they may also regulate distinct cellular functions via interactions at the membrane. Given the established role of Kv1.3 in both T-cells (adaptive immune cells) and microglia (innate immune cells of the brain), we examined the Kv1.3 interactome in these two cell types to identify cell type-specific, as well as conserved, interactions and associated pathways. We conducted a comparative analysis of the Kv1.3 interactome in JTCs with our previously published Kv1.3 interactome in BV-2 microglia (35). In our analysis, we identified a total of 1,845 Kv1.3 interactors in the JTC Kv1.3 AP dataset **(Fig. 2B and Fig. 4A)**. Applying the same threshold in BV-2 dataset, we found 863 Kv1.3 interactors **(Fig. 4A)**. Of these, 505 proteins were common Kv1.3 interactors across both cell lines **(Fig. 4B)**. We also found 1,340 proteins unique to JTCs and 358 unique to BV-2 cells, highlighting clear cell-type differences in the Kv1.3 channel proximity interactome **(Fig. 4B).** GO analyses of the enriched proteins indicated that the overlapping interactors are involved in Golgi vesicle transport (*e.g.* SCAMP1, CORO7, YKT6) and intracellular protein transport (*e.g.* TRAM1, TMED9, AHCYL1). The molecular functions associated with these overlapping proteins include cadherin binding (*e.g.* PTPN1, AHSA1, NOP56), cell adhesion molecule binding (*e.g.* TRIM25, DDX3X, FLNA) and translation regulator activity (*e.g.* GSPT1, GCN1, EIF4GI), with interactors primarily located in ER subcompartment (*e.g.* COPB4, LCLAT1, SEC16A), ER membrane (*e.g.* COPA, SGPL1, STIM1) and Golgi apparatus (*e.g.* PLIN3, GANAB, ARFGEF2) **(Fig. 4C)**. The presence of a high number of ER and Golgi proteins across cells may again be due to transient interactions during processing and trafficking of the Kv1.3 channel protein. Furthermore, the biological processes and molecular functions of the Kv1.3 interactome specific to JTCs appear to regulate GTPase regulator activity (*e.g.* RGS14, ARHGAP15, RAPGEF6), enzyme activator activity (*e.g.* PSME2, SMAP2, RPTOR) and protein phosphorylation (*e.g.* MAPKAP1, GRK3, PEAK1) **(Fig. 4C)**. In contrast, the distinct Kv1.3 interacting proteins in BV-2 cells are predominantly associated with transmembrane transport (*e.g.* ATP6V1C1, VMP1, TOMM22), oxidoreductase activity (*e.g.* COX4I1, NCF2, NDUFA9) and lipid biosynthetic process (*e.g.* ERLIN2, DGAT1, CLN6) **(Fig. 4C)**. In summary, Kv1.3 channels in T-cells appear to regulate cell activation, trafficking and signaling while in microglia primarily associate with proteins involved in mitochondrial function and lipid metabolism. Our analysis provides a molecular basis for functional variation of Kv1.3 channel in two distinct immune cell types.

**Figure 4:**
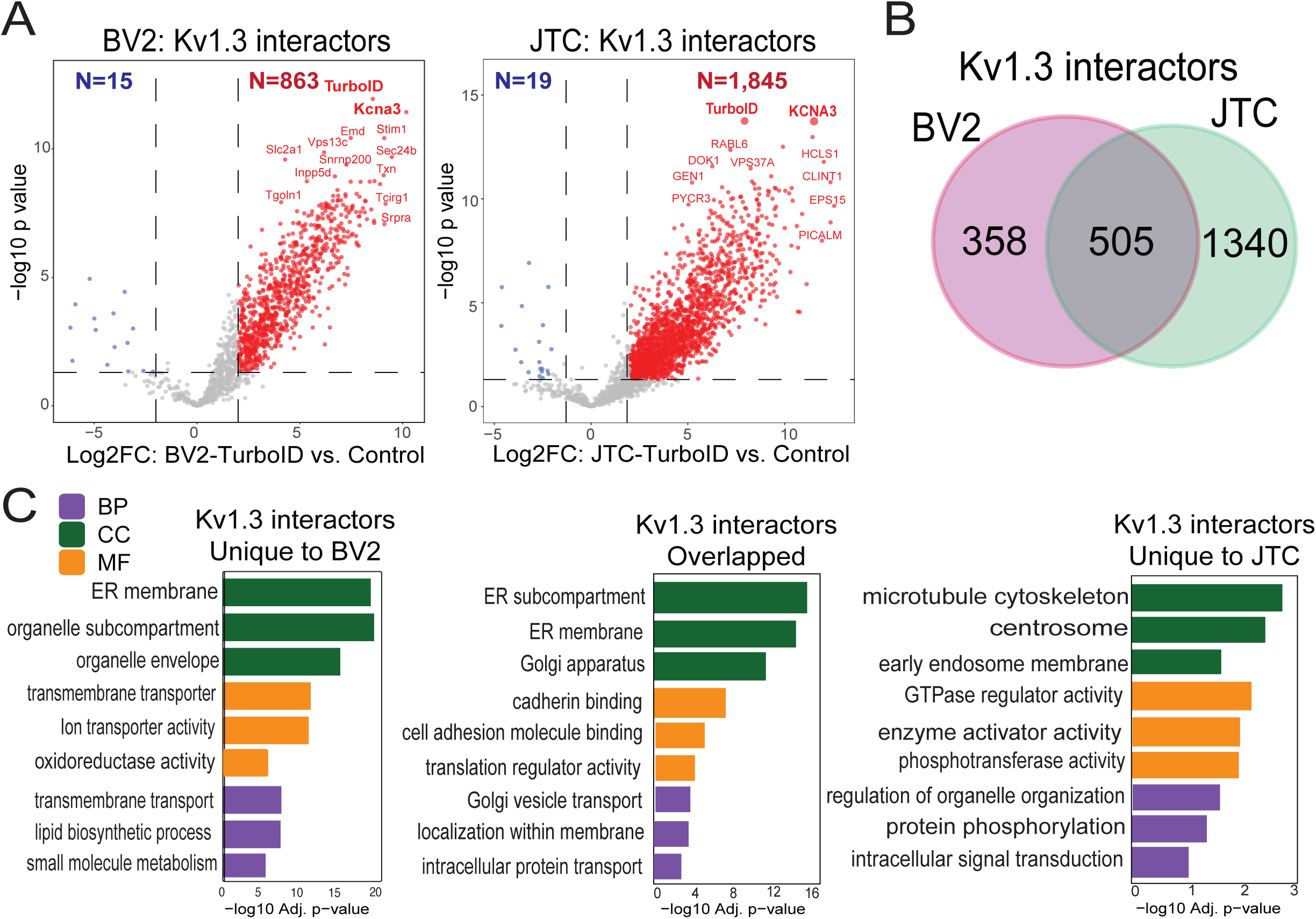
Comparative analysis of Kv1.3 channel interactome in BV-2 microglia and Jurkat T-Cells (JTC). **(A)** DEA between TurboID and Control sample identifies 863 interactors of Kv1.3 channel in BV-2 microglial cells while 1845 are interactors of Kv1.3 channel in JTC (n=3). **(B)** Intersection of BV-2 and JTC Kv1.3 interactors shows 358 and 1340 proteins unique to BV-2 and JTCs, respectively, along with an overlap of 505 channel interactors across given cell-types. **(C)** GO term enrichment analysis shows mitochondrial and metabolic function of Kv1.3 interactors in BV-2, while GTPase regulator and signaling role of channel interactors in JTCs. Overlapping proteins are associated with protein transport and localization. Detail analysis is given in Supp. Datasheet 5.

### Transmembrane proteins interactors of Kv1.3 channel in T-cells and microglia

Functional Kv1.3 channels are predominantly localized to the plasma membrane and the mitochondrial membrane, where they regulate membrane potential and K^+^ flux (37). Therefore, membrane-associated Kv1.3 channel interactors are of particular interest, as they are likely to mediate the most functionally significant interactions of Kv1.3 channels in immune cells.

We integrated the Kv1.3 AP proteome data with a list of transmembrane proteins (EBI-GO_Membrane and Membranome databases (https://membranome.org/proteins and https://www.ebi.ac.uk/QuickGO/term/GO:0016020). Of 3,732 known transmembrane proteins and 1,845 Kv1.3 interactors in JTCs, we identified 335 membrane-associated Kv1.3 interactors in T-cells **(Fig 5A**). These included distinct groups of plasma membrane (cell surface) interactors, mitochondrial membrane interactors, ER/Golgi, organelle and vesicle related membrane interactors. 55 plasma membrane Kv1.3 interacting proteins were identified (**Fig. 5B**), including known Kv1.3 interactors like integrins ITGB1 and ITGB2 (60) along with ITGAL, and CD3 complex proteins (40) (CD3D, CD3E, CD3G). 32 proteins were associated with the mitochondrial membrane (**Fig. 5B**). Additionally, 18 proteins were identified as ER membrane components **(Supp. Fig. 4A**), along with several proteins attributed to Golgi membrane. We also identified 25 members of the SNAP receptor activity family (**Supp. Fig. 4B**), suggesting the potential involvement of SNAP receptor-mediated vesicle trafficking in the localization of Kv1.3 channels to the membrane. Alternatively, Kv1.3 channels may also influence SNARE proteins that regulate exocytosis, with functional roles in human effector T cells (61).

**Figure 5:**
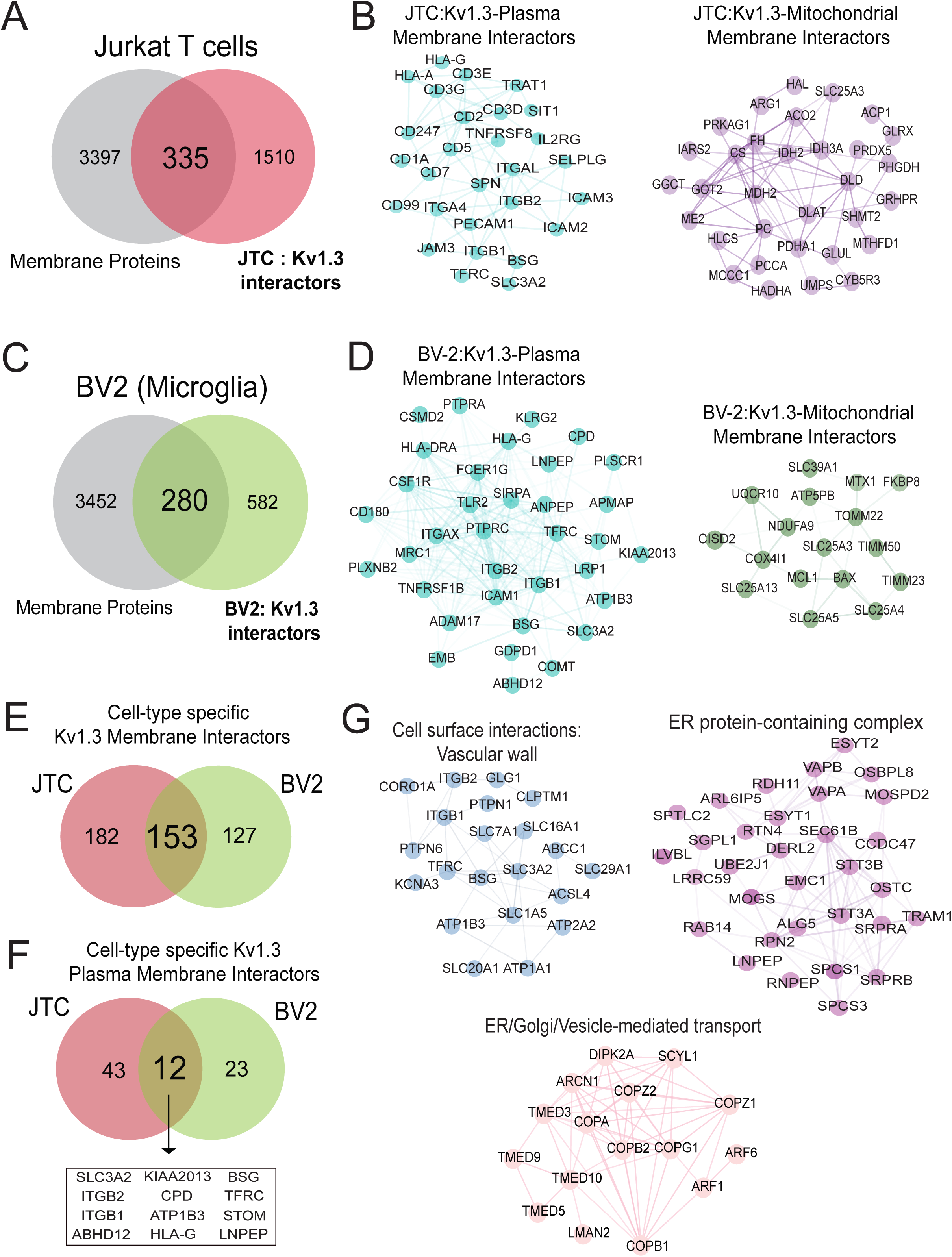
Transmembrane interactors of Kv1.3 channel in BV-2 microglia and Jurkat T-Cells (JTC). **(A)** Venn diagram of JTC-Kv1.3 interactors and membrane proteins highlight 335 membrane proteins in Kv1.3 interactome in JTCs. **(B)** STRING network analysis of 335 membrane interactors of Kv1.3 channel identifies 55 plasma membrane and 32 mitochondrial membrane proteins interacting with Kv1.3 channel in JTCs. **(C)** Venn diagram analysis between membrane proteins and BV-2 Kv1.3 interactors shows 280 membrane proteins in Kv1.3 interactome in BV-2 microglial cell. **(D)** Network analysis of 280 membrane Kv1.3 interactors in BV-2 illustrates 35 and 14 of these are plasma and mitochondrial membrane interactors, respectively. **(E)** Intersection between membrane interactors of Kv1.3 channel in BV-2 and JTCs shows 182 are JTC and 127 are BV-2 unique interactors, while 153 are shared across cell-type. **(F)** Intersection between plasma membrane interactors of Kv1.3 channel in BV-2 and JTC displays 43 distinct interactors in JTCs, while 23 are specific to BV-2 cells. 12 shared plasma membrane interactors among JTCs and BV-2 are highlighted. **(G)** STRING analysis reveals functional clustering of overlapping proteins (153) in JTC and BV-2 Kv1.3 membrane interactors intersection given in Fig 5E. Panels highlight proteins involved in cell surface interactions at the vascular wall, focusing on adhesion and signaling, proteins associated with ER to Golgi vesicle-mediated transport, emphasizing their roles in intracellular trafficking and vesicle formation and ER protein-containing complex, illustrating proteins critical for ER organization and protein folding. Complete table of each analysis is given in Supp. Datasheet 6.

We also contrasted membrane-associated Kv1.3 interactors in JTCs with BV-2 microglia, using a similar approach outlined above. 280 transmembrane proteins were present in the BV-2 Kv1.3 interactome (**Fig. 5C**), which included 35 plasma membrane (*e.g.* CD180, LRP1, CSF1R) **Fig. 5D**) and 14 mitochondrial membrane proteins (**Fig. 5D**). Mitochondrial membrane interactors in BV2 microglia included important inner (TIMM23, TIMM50) and outer membrane (TOMM22, MTX1) proteins, which were not present in the JTC interactome. 153 transmembrane interactors of Kv1.3 were shared across JTCs and BV2 microglia while 182 were unique to JTCs and 127 were unique to BV-2 microglia (**Fig. 5E**). 12 shared plasma membrane interactors were identified, including ITGB1, ITGB2, SLC3A2, STOM (**Fig. 5F**)., while 43 and 23 were specific to JTC and BV-2, respectively. In contrast to the partial overlap of plasma membrane Kv1.3 interactors in microglia and T-cells, mitochondrial membrane Kv1.3 interactors across cell types showed no overlap.

Overlapping membrane interactors of JTCs and BV-2 include solute transporters (SLCs) and integrins that mediate cell surface interactions at the vascular wall, proteins from the TMED and COP family, enriching ER to Golgi vesicle-mediated transport pathway, and proteins belonging to ER protein-containing complexes like SPCS1, STT3A, RAB14 (**Fig. 5G**). Overall, Kv1.3’s transmembrane interactome in JTCs and BV-2 cells included both shared and unique interactors that contribute to its diverse roles. The overlap of 153 interactors between JTCs and BV-2 cells points to conserved mechanisms of Kv1.3 regulation and function, while the unique interactors reflect cell-type-specific adaptations that likely contribute to the distinct physiological roles of Kv1.3 in T-cells and microglia.

### Nomination of Kv1.3 channel interactors with causal roles in T cell-mediated autoimmune diseases

Activated TEMs highly express Kv1.3 channels and are implicated in the pathogenesis of autoimmune diseases (23). To nominate Kv1.3 interacting proteins that are most relevant to autoimmune diseases, we obtained a list of 2,327 autoimmune disease risk genes from the NHGRI-EBI GWAS catalog for various autoimmune diseases (https://www.ebi.ac.uk/gwas/efotraits/EFO_0005140). Of these, 178 proteins were identified as Kv1.3 channel interacting proteins in JTCs **(Fig 6A)**. Our analysis highlighted interaction of Kv1.3 channel with genes, such as UBASH3A, ITSN2, ERAP1 and EIF4H, linked to multiple autoimmune conditions including type-1 diabetes (T1D), rheumatoid arthritis (RA), multiple sclerosis (MS), systemic lupus erythematosus (SLE), inflammatory bowel disease (IBD), and celiac disease, which were highly represented in the Kv1.3 interactome. Additionally, other abundant proteins in the AP proteome included risk genes associated with IBD (*e.g*. PRRC2A, PTK2B, SLAIN2, ALMS1), RA (*e.g*. LZTS1, PRRC2A, PRR12), Crohn’s disease (*e.g*. PTK2B, ERGIC1, SLAIN2), SLE (*e.g.* PRR12, RABGAP1L, MAPT) and ulcerative colitis (ERGIC1). GO analyses showed enrichment of adaptive immune response/immune regulation, cell adhesion, signal transduction, binding signaling receptors and kinases **(Fig 6B).** STRING analysis highlighted TCR complex (UBASH3, PTPN22, PTPN1, CD3E) **(Fig 6C)**, cytokine-mediated signaling (IRAK1, RUNX1, CLEC16A, STIM1) **(Fig 6D)**, and actin filament bundle assembly/lymphocyte migration (ITGA4, SENP1, NFKBIE, ARHGAP4) **(Fig 6E)** proteins as autoimmune disease-relevant Kv1.3 interactors. These Kv1.3 interactions provide a framework for understanding how Kv1.3 channels may regulate immune mechanisms in autoimmune diseases and guide target nomination for validation studies.

**Figure 6:**
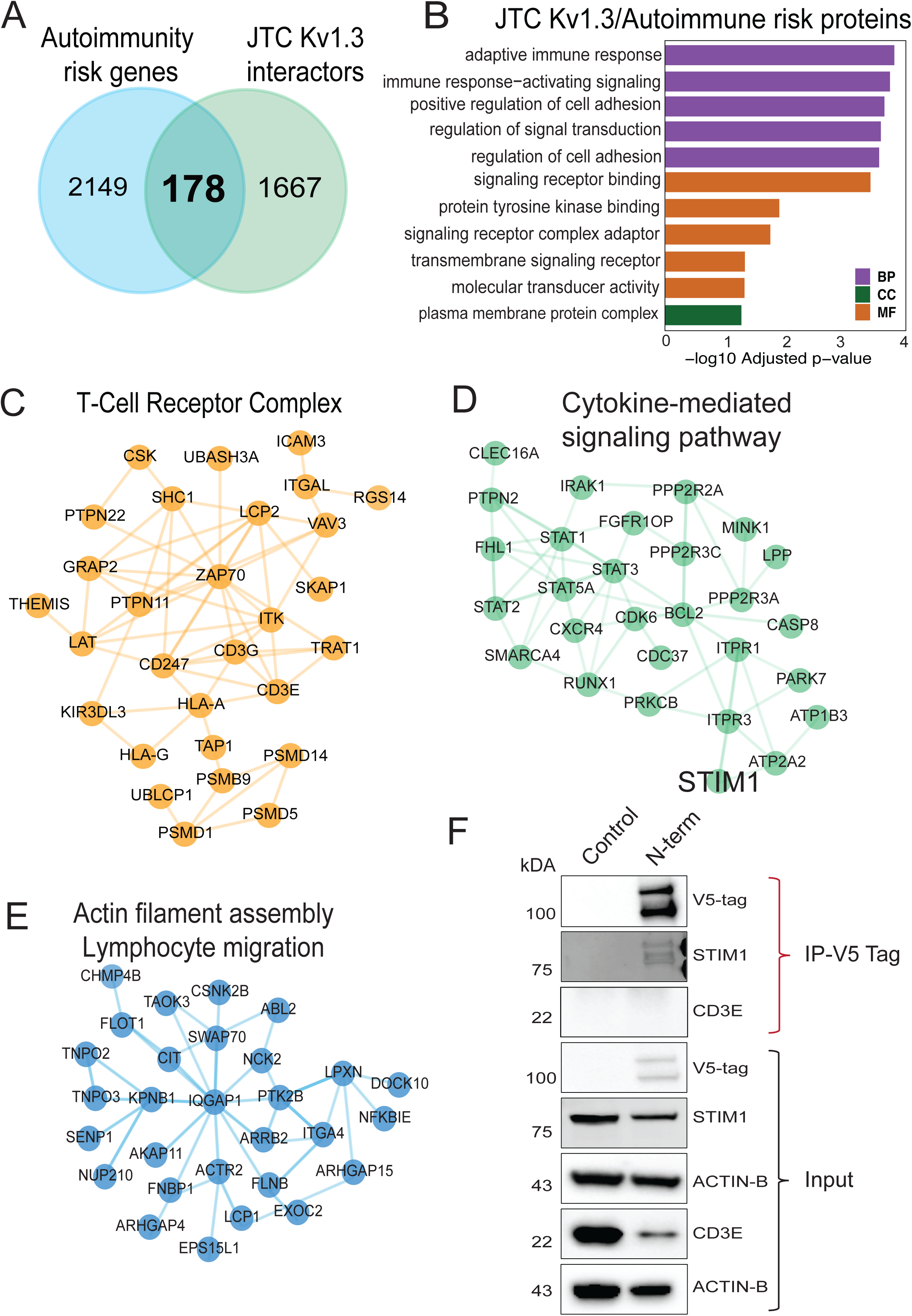
Comparative molecular analyses between Kv1.3 interactors and autoimmune diseases associated risk genes. **(A)** Intersection of autoimmune diseases associated risk genes and Kv1.3 interactors shows 178 Kv1.3 enriched proteins in JTCs are risk factor for autoimmune conditions. **(B)** GO term enrichment analysis of 178 Kv1.3 interacting autoimmune risk genes highlights their role in immune response and signaling. STRING analyses illustrating protein-protein interactions shows **(C)** T-cell receptor complex, **(D)** Cytokine-mediated signaling pathway and **(E)** Actin filament bundle assembly or lymphocyte migration functional clusters in Kv1.3 interacting autoimmune risk genes found in JTCs. **(F)** Western blot following co-immunoprecipitation showing Kv1.3 direct interaction with STIM1 and no physical interaction with CD3E, using V5-tag fused to TurboID. (IP-Immunoprecipitation) (n=3). Table showing details of each analysis is given in Supp. Datasheet 7.

Based on strength of enrichment in the Kv1.3 interactome and strength of genetic risk association, we nominated STIM1 and CD3E for validation studies of physical interaction with Kv1.3 using co-immunoprecipitation. Calcium entry with the STIM-ORAI (CRAC) channel is a critical regulator of T cell activation and effector functions. STIM1 is also a target for immune modulation (62). Our data from JTCs nominated STIM1 as a top Kv1.3 proximity protein (log2 fold change >9). We also selected CD3E (log2 fold change=5), a part of the TCR-CD3 complex that plays a central role in T-cell activation and immune response (63). CD3 and Kv1.3 molecular proximity is well-known (40, 64).To validate these interactions, we leveraged the V5 tag in our V5-TurboID-Kv1.3 fusion construct to perform V5 tag antibody-based co-immunoprecipitation experiments to enrich physical interactors of Kv1.3 channel proteins. Kv1.3 and STIM1 were found to co-precipitate, confirming a direct physical protein-protein interaction between two proteins in JTCs **(Fig. 6F).** This highlights a functional relationship between Kv1.3 and STIM1, which is particularly relevant in our cell model, where both Kv1.3 and STIM1 are critical for T-cell activation, proliferation, and cytokine production (62, 65, 66). Conversely, CD3E band was not detectable upon co-immunoprecipitation suggesting no physical interaction with Kv1.3, implying that the CD3 complex is within proximity of the Kv1.3 channel but not directly interacting with it **(Fig. 6F)**. This reinforces earlier known Kv1.3 channel interactions with the TCR-CD3 complex through intermediatory proteins (67).

## DISCUSSION

Kv1.3 channels modulate activation, proliferation and cytokine production in T-cells (65, 66). Although Kv1.3 has been well studied in T-cells, understanding its precise molecular functions and signaling roles deserve further investigation. We used a bioengineered enzyme, TurboID, fused to the Kv1.3 protein to label and identify proteins within proximity of the channel in the JTC human T-cell line (42). We generated JTC lines, stably-expressing Kv1.3 channels fused to TurboID at either N-terminal, C-terminal or C-terminal⊗PDZ, and validated functional transduction of channel through electrophysiological studies. We excluded the effect of transduction on basal T-cell activation and functions by analyzing the effect of channel overexpression and blockade on NFAT signaling, certain kinases and cytokines. After MS-based quantification of biotinylated proteins, we identified over 1,800 proteins as interactors of Kv1.3, which included several previously known protein interactors such as integrins (ITGB1, ITGB2), immune synapse proteins (DLG4, DLG1), apoptosis-related mitochondrial protein (BCL-2), regulatory subunits (KCNAB2) and immune signaling proteins (STAT1, STAT3), providing validation of our approach. In addition to confirming these known interactors, our study provides a surprisingly broad network of proteins in proximity to Kv1.3 channels in T-cells that likely includes indirect, transient, and stable interactors. Several Kv1.3 interactors also showed preferential interaction with the N-terminus and with the C-terminus of the channels, with a small, yet important, subsubset of C-terminal interactors showing dependency on the PDZ-binding domain. While many Kv1.3-interacting proteins were identified in both T-cells and microglia, several are unique to each cell type. Although Kv1.3 primarily regulate K^+^ efflux, membrane potential and Ca^2+^ influx as a general mechanism of immune regulation, the interactome of Kv1.3 clearly varies across immune cell types, suggesting that it may also regulate a plethora of different immune functions (35, 36). Importantly, the Kv1.3 interactome identified by our proximity labeling approach revealed 178 proteins with causal roles in the pathogenesis of autoimmune diseases, highlighting that Kv1.3 channels interact with and potentially modulate these disease-relevant pathways.

Two recent publications, one by Bowen *et al*. from our group and another by Prosdocimi *et al*., have employed proximity-dependent labeling techniques to identify the domain-specific interactomes of Kv1.3 channels in BV-2 and HEK293 cells (35, 36). In our analysis, we found that proteins that interact with the N-terminus of Kv1.3 are involved in protein synthesis and trafficking, Golgi vesicle transport, vesicle-mediated transport, mRNA processing and endosomal processes which are in alignment with previous reports (35). A noteworthy N-terminal interaction identified in this study is with the immune adaptor protein STING1. STING1 triggers production of type 1 interferons and initiates a cascade of events involving immune cell activation and inflammation (68). Further investigations are required to find out the functional implication of this interaction. The interaction of the Kv1.3 C-terminus with proteins implicated involved in cell junction maintenance, immune signaling, and immune synapse organization are consistent with previous findings indicating the critical role of Kv1.3 channel in T-cell activation and immune synapse formation (69). Small GTPases are essential for T-cell activation, migration, and immune synapse formation (70, 71). Though there are some studies suggesting an association between Kv1.3 channels and GTPase activity, the exact regulatory role of Kv1.3 remains unclear. Similar to our observations, Prosdocimi *et al*. found that Kv1.3-enriched fractions are involved in GTPase-related functions (36). Our data indicated that C-terminus-specific interactions are particularly enriched in proteins associated with GTPase activity and regulation, with certain interactions mediated through its PDZ-binding domain. Additionally, the C-terminus interacts with TSC1, a negative regulator of critical cellular growth, metabolic pathways, and mTOR signaling (72), suggesting Kv1.3 may act as a regulatory unit in cell metabolism. This interaction may explain the higher metabolic rate observed in Kv1.3 KO mice (73). Interestingly, the Kv1.3 channel C-terminus also interacts with LAMTOR1, an activator of the mTOR pathway (74). In physiological conditions, it would be interesting to explore how these interactions are balanced within the cell to ensure proper cellular function. Although the N-terminal interactome of Kv1.3 appears to be larger than the C-terminal interactome, it may capture many protein processing and trafficking proteins that likely reflect interactions that occur in the ER/Golgi and during trafficking to the cell surface. Because the C-terminal interactome includes signaling and GTPase related proteins, it is likely that C-terminus of Kv1.3 may regulate critical immune functions. STAT1 was recently identified as a C-terminal interactor of Kv1.3 and indeed, Kv1.3 blockade in microglia reduced STAT1 phosphorylation (35). In a melanoma cell line, Kv1.3 interactions with STAT3 protein were also verified, linking Kv1.3 channel function with immune signaling (36).

According to Bowen *et al.,* in BV-2 microglia, deletion of the PDZ-binding domain had significant impact on the C-terminal interactome (35). Kv1.3 channel loses 70 interactions upon deletion of this domain, consisting of proteins associated with crucial processes like antigen processing and presentation and immune signaling pathways. However, our study in JTCs showed modest effects of deletion of the C-terminal PDZ-binding TDV motif on the Kv1.3 interactome. The channel lost interactions with five proteins, including ROCK2, a serine/threonine kinase that plays a significant role in enhancing the pathogenic functions of T-cells. ROCK2 is also associated with T cells activation in IBD (75). Another notable PDZ-binding domain dependent interaction of Kv1.3 channel is with TUBB6, a beta-tubulin isoform that forms microtubules. Co-expression analysis of TUBB6 highlights its correlation with T-cell proliferation in immune signaling and CD8+ T-cells infiltration in carcinomas (76). Additionally, the shift observed in C-terminus interactions, with domain deletion, implies that the loss of the PDZ-binding domain not only weakens specific regulatory interactions but may also recruit non-specific or compensatory interactions that could alter Kv1.3 channel functionality. One of the notable interactions of Kv1.3 channel upon PDZ-binding domain deletion is with FYN kinase, a neuroinflammation modulator and therapeutic target in neurodegenerative diseases like Alzheimer’s and Parkinson diseases (77, 78). Previously, Sarkar *et al.* have shown that FYN binds to Kv1.3 and regulates channel activity in microglia (79). Interestingly, in our analysis FYN interaction gets stronger with PDZ-binding domain deletion, suggesting that this domain has an inhibitory effect on FYN-Kv1.3 interaction. However, how this interaction relates to FYN or Kv1.3 channel function warrants further investigation. Moreover, the gain of ER-associated interactors suggests that PDZ-binding domain deletion may alter channel localization or turnover, potentially impacting Kv1.3-mediated signaling dynamics in T-cells. Our findings of PDZ-binding domain interactions and the observed changes with PDZ-binding domain deletion further emphasize the role of the Kv1.3 channel C-terminus in the structural organization of immune signaling complexes and modulating T-cell function.

An important observation in our study is that nearly 500 Kv1.3 interactors are shared across two major immune cell lines (T cells and microglia), both of which express Kv1.3 channels (27, 80). These common interactors are related to protein transport and localization, along with cell signaling functions like cadherin and cell adhesion molecule binding. However, we found that the majority (∼73%) of Kv1.3 interactors in JTCs are unique to T-cells, while 42% of Kv1.3 interactors in BV-2 microglia are unique to microglia. These represent cell type-specific Kv1.3 interactors, providing a framework for how different cell types may link Kv1.3 channels to distinct molecular machinery and functional pathways. Interestingly, Kv1.3 interactors in T cells are predominantly associated with regulating cell signaling. While those in microglia are involved in cellular metabolism and energy production. The Kv1.3 channel in T-cells interacts with various kinases, including IRAK1, CDK1, STK26, ROCK2, and MARK3, highlighting its role in protein phosphorylation. This involvement in protein modifications might contribute to the channel’s immunomodulatory function. These differences suggest that Kv1.3 may differentially regulate cell activation and signaling functions in T-cells compared to metabolic function in microglia.

Furthermore, our analyses of plasma membrane Kv1.3 interactors helped us identify important cell surface interactions. In T-cells, we found that Kv1.3 interacts with TCR complex proteins such as CD3 components and CD247, which are essential for T-cell activation and antigen recognition (81). In addition, we found receptor kinases PTPRK, PTP7, as well as cytokine signaling proteins (*e.g.* IL2RG), essential for lymphocyte proliferation and immune regulation (82). Similarly, in microglia we found that the receptor tyrosine kinase CSF1R is in proximity to Kv1.3 channels. CSF1R regulates macrophage and microglial development, survival, differentiation and proliferation with important roles in neurological disease pathogenesis (83). Proteins like SIRPA, LRP1, that have a role in phagocytosis were also identified as microglia-specific interactors of Kv1.3 (84, 85). The shared interactors in T-cells and microglia underscore fundamental roles of Kv1.3 that span both immune and glial cell types. Conversely, the cell-type-specific interactors recognize specialized functions tailored to immune responses in T-cells and neuroinflammatory regulation in microglia. These findings provide a foundation and framework for future mechanistic studies to investigate the functional relevance of these interactions, particularly in the context of Kv1.3’s roles in immunity, neuroinflammation, and cell metabolism. However, it is important to note that our analyses were conducted using human T-cell and mouse microglial lines. While we expect most interactors to be the same, some may not be fully consistent across species.

Our study also identified several Kv1.3 interactors that are localized to mitochondria. Kv1.3 channels have been shown to be present in the inner mitochondrial membrane and may regulate mitochondrial function and apoptosis (86). We indeed identified BCL-2, a direct interactor of the known mitochondrial Kv1.3 interactor, BAX (38). In T-cells, the mitochondrial Kv1.3 interactors primarily include proteins such as ACO2, CS, FH, IDH2, and IDH3A, which are functionally related to energy production and mitochondrial metabolism, as well as GOT2, GLUL, SHMT2, and PHGDH, which are associated with amino acid metabolism. In microglia, Kv1.3 interactors in the mitochondria included proteins such as TIMM50, TOMM22, and MTX1, which are involved in protein import and maintaining mitochondrial integrity (87–89). There are also several critical components of the mitochondrial electron transport chain (ETC) and ATP synthase (Complex V) (UQCR10, COX4I1, ATP5PB, and ATP5MK), present in the microglial interactome, which enable oxidative phosphorylation and drive ATP production (90–92). Surprisingly, Kv1.3 interactors in the mitochondria were not shared across microglia and T-cells. These distinctions might be due to variations in the functions of mitochondria in microglia, where they manage the high energy demand required for brain immunosurveillance (93), in contrast to T-cells, where mitochondrial metabolism is primarily important for T-cell activation and proliferation (94). While our studies did not obtain Kv1.3 interactions from mitochondria-enriched cellular fractions, our results are in agreement with mitochondrial localization and functions of Kv1.3 channels.

Kv1.3 is highly expressed by autoreactive activated TEMs in several autoimmune diseases (23). Kv1.3 channel blockers, such as PAP-1 (5-(4-phenoxybutoxy) psoralen) and sea anemone peptides (ShK-186, ShK-223) are effective in suppressing autoimmune responses and improving neurological outcomes in rodent models of MS (23), T1D (20), arthritis (20), psoriasis (95), autoimmune encephalomyelitis (96), granulomatosis with polyangiitis (97) and systemic lupus erythematosus (98). To gain a better understanding of how Kv1.3 channels regulate immune functions across this spectrum of autoimmune diseases, we identified several autoimmune disease risk gene products as Kv1.3 interactors. Among these, STIM1 was identified and validated as an interacting partner of Kv1.3 using co-immunoprecipitation. While the functional relationship between Kv1.3 channels and the CRAC channel components has been observed, along with localization of STIM1, ORAI and Kv1.3 at immunological synapses during T-cell activation, the direct evidence of physical interaction between Kv1.3 and STIM1 is lacking (41, 99, 100). Our results therefore provide another direct link between Kv1.3 channel function and the CRAC channel in T-cells. In addition, CD3E, another identified protein, was validated only to be in proximity of the channel, but not physically interacting with it. This highlights the role of adaptor proteins in mediating the interaction of Kv1.3 channels with the TCR-CD3 complex at immunological synapses (67). Since aberrant abnormalities of T-cell activation and cytokine signaling are hallmarks of autoimmune diseases, our findings provide important molecular links between Kv1.3 channel function and disease-relevant immune mechanisms.

We acknowledge several limitations of our study. Firstly, although JTCs are a well-established model for studying T-cells *in vitro,* these immortalized and permanently activated cells, lacking CD4 and CD8 surface receptors, may not fully recapitulate the signaling and functional characteristics of primary T cells *in vivo*. Further validation of interactions in primary T-cells or in an *in vivo* model are needed. Secondly, proximity labeling using TurboID may capture proteins in the vicinity of Kv1.3 that do not directly interact with the channel, and may represent transient or less stable interactions, not all of which may be functionally important. As we did for STIM1, additional validation with co-immunoprecipitation should therefore be considered for targets of interest. Lastly, the comparative analysis of Kv1.3 channel functions between JTCs and BV-2 cells may be partly limited by inter-species differences, as JTCs are human cells while BV-2 cels are a mouse line.

Overall, this study highlights the possible role of Kv1.3 channels in various immune and signaling pathways. Future research should focus on validating and elucidating the functional consequences of the identified Kv1.3 channel interactions. These analyses should be extended to other immune cells, with varied cellular activation states and disease contexts. It is also highly important to validate key interactions in primary T-cells and in *in-vivo* models. Future studies testing the effect of channel blockade on these nominated pathways will deepen our understanding of the role of Kv1.3 in T-cell signaling and autoimmune diseases and its potential as a therapeutic target.

## CONCLUSIONS

In conclusion, our study showcases how high-throughput interactomics, target nomination and focused validation can be used to identify novel protein-protein interactors of an ion channel in immune cells. Using proximity-based proteomics in two mammalian cell lines, we identified a large network of Kv1.3 channel interacting proteins, some of which validate prior findings while the majority represent novel interactions, including physical interactions between Kv1.3 and the CRAC channel protein STIM1. Our findings align with earlier research suggesting that the potassium ion channel Kv1.3 contributes to signaling cascades necessary for immune responses. The C-terminus directs these immune interactions, with several significant interactions mediated by the PDZ-binding domain, while the N-terminus regulates protein synthesis, trafficking and localization. We have highlighted many membrane specific interactors of Kv1.3 channel in T-cells and distinguished them from microglial specific interactors. Lastly, we provide additional evidence linking Kv1.3 channels with mechanisms of T cell-mediated autoimmunity and suggest a framework for future mechanistic investigations to further explore Kv1.3 channel as a therapeutic target for autoimmune diseases.

## Supporting information

Supplementary Datasheet 1

Supplementary Datasheet 2

Supplementary Datasheet 3

Supplementary Datasheet 4

Supplementary Datasheet 5

Supplementary Datasheet 6

Supplementary Datasheet 7

## ABBREVIATIONS

TCR: T-cell receptor
APC: Antigen presenting cells
TEM: T effector memory cels
TCM: T central memory cells
IGF: Insulin-like growth factor
EGF: Epidermal growth factor
LFQ-MS: Label-free quantitative mass spectrometry
DE: Differential enrichment analysis
DEP: Differentially expressed proteins

## STATEMENTS

## Acknowledgements

This research was supported by the Viral Vector Core of the Emory Center for Neurodegenerative Disease Core Facilities, the Emory Flow Cytometry Core (EFCC), the Custom Cloning Core Division of Emory Integrated Genomics Core (EIGC), and the Integrated Cellular Imaging Core (ICI) all at Emory University. BioRender for software to create images.

## Author’s contributions

Conceptualization: D.K., C.A.B., N.T.S., H.W., S.R.

Methodology: D.K., C.A.B., U.S., H.M.N., R.K., P.K., A.D.B., S.B., B.R.T., L.B.W., N.T.S., H.W., S.R.

Investigation: D.K., C.A.B., U.S., N.T.S., S.R.

Writing-Original draft: D.K., C.A.B., S.R.

Writing-Review and Editing: D.K., C.A.B., U.S., H.M.N., R.K., P.K., A.D.B., S.B., B.R.T., L.B.W., N.T.S., H.W., S.R.

Funding acquisition: C.A.B, L.B.W, N.T.S., S.R.

Resources: N.T.S., S.R.

Supervision: N.T.S., S.R.

## Statement of Ethics

The authors have no ethical conflicts to disclose.

## REFERENCES

1. Shah K, Al-Haidari A, Sun J, Kazi JU. T cell receptor (TCR) signaling in health and disease. Signal transduction and targeted therapy. 2021;6(1):412.

2. Hosokawa H, Rothenberg EV. Cytokines, transcription factors, and the initiation of T-cell development. Cold Spring Harbor perspectives in biology. 2018;10(5):a028621.

3. Hosokawa H, Rothenberg EV. How transcription factors drive choice of the T cell fate. Nature Reviews Immunology. 2021;21(3):162–76.

4. Sun L, Su Y, Jiao A, Wang X, Zhang B. T cells in health and disease. Signal transduction and targeted therapy. 2023;8(1):235.

5. He X, He X, Dave VP, Zhang Y, Hua X, Nicolas E, et al. The zinc finger transcription factor Th-POK regulates CD4 versus CD8 T-cell lineage commitment. Nature. 2005;433(7028):826–33.

6. Yu Q, Erman B, Bhandoola A, Sharrow SO, Singer A. In vitro evidence that cytokine receptor signals are required for differentiation of double positive thymocytes into functionally mature CD8+ T cells. The Journal of experimental medicine. 2003;197(4):475–87.

7. Topchyan P, Lin S, Cui W. The role of CD4 T cell help in CD8 T cell differentiation and function during chronic infection and cancer. Immune Network. 2023;23(5).

8. Laidlaw BJ, Craft JE, Kaech SM. The multifaceted role of CD4+ T cells in CD8+ T cell memory. Nature Reviews Immunology. 2016;16(2):102–11.

9. Koh C-H, Lee S, Kwak M, Kim B-S, Chung Y. CD8 T-cell subsets: heterogeneity, functions, and therapeutic potential. Experimental & Molecular Medicine. 2023;55(11):2287–99.

10. Kaech SM, Tan JT, Wherry EJ, Konieczny BT, Surh CD, Ahmed R. Selective expression of the interleukin 7 receptor identifies effector CD8 T cells that give rise to long-lived memory cells. Nature immunology. 2003;4(12):1191–8.

11. Joshi NS, Cui W, Chandele A, Lee HK, Urso DR, Hagman J, et al. Inflammation directs memory precursor and short-lived effector CD8+ T cell fates via the graded expression of T-bet transcription factor. Immunity. 2007;27(2):281–95.

12. Rangaraju S, Chi V, Pennington MW, Chandy KG. Kv1. 3 potassium channels as a therapeutic target in multiple sclerosis. Expert opinion on therapeutic targets. 2009;13(8):909–24.

13. Samakai E, Go C, Soboloff J. Defining the roles of Ca2+ signals during T cell activation. Signaling Mechanisms Regulating T Cell Diversity and Function. 2017:177–202.

14. Carreras-Sureda A, Zhang X, Laubry L, Brunetti J, Koenig S, Wang X, et al. The ER stress sensor IRE1 interacts with STIM1 to promote store-operated calcium entry, T cell activation, and muscular differentiation. Cell Reports. 2023;42(12).

15. Feske S, Wulff H, Skolnik EY. Ion channels in innate and adaptive immunity. Annual review of immunology. 2015;33(1):291–353.

16. Bose T, Cieślar-Pobuda A, Wiechec E. Role of ion channels in regulating Ca2+ homeostasis during the interplay between immune and cancer cells. Cell death & disease. 2015;6(2):e1648-e.

17. Arango M-T, Shoenfeld Y, Cervera R, Anaya J-M. Infection and autoimmune diseases. Autoimmunity: from bench to bedside [Internet]: El Rosario University Press; 2013.

18. Ye Z, Shen Y, Jin K, Qiu J, Hu B, Jadhav RR, et al. Arachidonic acid-regulated calcium signaling in T cells from patients with rheumatoid arthritis promotes synovial inflammation. Nat Commun. 2021;12(1):907.

19. Park YJ, Yoo SA, Kim M, Kim WU. The Role of Calcium-Calcineurin-NFAT Signaling Pathway in Health and Autoimmune Diseases. Front Immunol. 2020;11:195.

20. Beeton C, Wulff H, Standifer NE, Azam P, Mullen KM, Pennington MW, et al. Kv1. 3 channels are a therapeutic target for T cell-mediated autoimmune diseases. Proceedings of the National Academy of Sciences. 2006;103(46):17414–9.

21. Birkner K, Wasser B, Ruck T, Thalman C, Luchtman D, Pape K, et al. β1-Integrin–and K V 1.3 channel–dependent signaling stimulates glutamate release from Th17 cells. The Journal of clinical investigation. 2020;130(2):715–32.

22. Ohya S, Matsui M, Kajikuri J, Endo K, Kito H. Increased interleukin-10 expression by the inhibition of Ca2+-activated K+ channel KCa3. 1 in CD4+ CD25+ regulatory T cells in the recovery phase in an Inflammatory Bowel Disease Mouse Model. Journal of Pharmacology and Experimental Therapeutics. 2021;377(1):75–85.

23. Wulff H, Calabresi PA, Allie R, Yun S, Pennington M, Beeton C, et al. The voltage-gated Kv1. 3 K+ channel in effector memory T cells as new target for MS. The Journal of clinical investigation. 2003;111(11):1703–13.

24. Gocke AR, Lebson LA, Grishkan IV, Hu L, Nguyen HM, Whartenby KA, et al. Kv1. 3 deletion biases T cells toward an immunoregulatory phenotype and renders mice resistant to autoimmune encephalomyelitis. The Journal of Immunology. 2012;188(12):5877–86.

25. Matteson D, Deutsch C. K channels in T lymphocytes: a patch clamp study using monoclonal antibody adhesion. Nature. 1984;307(5950):468-71.

26. DeCoursey TE, Chandy KG, Gupta S, Cahalan MD. Voltage-gated K+ channels in human T lymphocytes: a role in mitogenesis? Nature. 1984;307(5950):465-8.

27. Selvakumar P, Fernández-Mariño AI, Khanra N, He C, Paquette AJ, Wang B, et al. Structures of the T cell potassium channel Kv1. 3 with immunoglobulin modulators. Nature Communications. 2022;13(1):3854.

28. Nicolaou SA, Szigligeti P, Neumeier L, Molleran Lee S, Duncan HJ, Kant SK, et al. Altered dynamics of Kv1. 3 channel compartmentalization in the immunological synapse in systemic lupus erythematosus. The Journal of Immunology. 2007;179(1):346–56.

29. Comes N, Bielanska J, Vallejo-Gracia A, Serrano-Albarrás A, Marruecos L, Gómez D, et al. The voltage-dependent K+ channels Kv1. 3 and Kv1. 5 in human cancer. Frontiers in physiology. 2013;4:283.

30. Doyle DA, Cabral JM, Pfuetzner RA, Kuo A, Gulbis JM, Cohen SL, et al. The structure of the potassium channel: molecular basis of K+ conduction and selectivity. science. 1998;280(5360):69–77.

31. Pérez-García MT, Cidad P, López-López JR. The secret life of ion channels: Kv1. 3 potassium channels and proliferation. American Journal of Physiology-Cell Physiology. 2018;314(1):C27–C42.

32. Navarro-Pérez M, Estadella I, Benavente-Garcia A, Orellana-Fernández R, Petit A, Ferreres JC, et al. The Phosphorylation of Kv1. 3: A Modulatory Mechanism for a Multifunctional Ion Channel. Cancers. 2023;15(10):2716.

33. Gamper N, Fillon S, Huber S, Feng Y, Kobayashi T, Cohen P, et al. IGF-1 up-regulates K+ channels via PI3-kinase, PDK1 and SGK1. Pflügers Archiv-European Journal of Physiology. 2002;443:625–34.

34. Martínez-Mármol R, Comes N, Styrczewska K, Pérez-Verdaguer M, Vicente R, Pujadas L, et al. Unconventional EGF-induced ERK1/2-mediated Kv1. 3 endocytosis. Cellular and Molecular Life Sciences. 2016;73:1515–28.

35. Bowen CA, Nguyen HM, Lin Y, Bagchi P, Natu A, Espinosa-Garcia C, et al. Proximity Labeling Proteomics Reveals Kv1.3 Potassium Channel Immune Interactors in Microglia. Mol Cell Proteomics. 2024;23(8):100809.

36. Prosdocimi E, Carpanese V, Todesca LM, Varanita T, Bachmann M, Festa M, et al. BioID-based intact cell interactome of the Kv1. 3 potassium channel identifies a Kv1. 3-STAT3-p53 cellular signaling pathway. Science Advances. 2024;10(36):eadn9361.

37. Capera J, Navarro-Pérez M, Moen AS, Szabó I, Felipe A. The mitochondrial routing of the Kv1. 3 channel. Frontiers in oncology. 2022;12:865686.

38. Szabó I, Bock J, Grassmé H, Soddemann M, Wilker B, Lang F, et al. Mitochondrial potassium channel Kv1. 3 mediates Bax-induced apoptosis in lymphocytes. Proceedings of the National Academy of Sciences. 2008;105(39):14861–6.

39. Jang SH, Byun JK, Jeon W-I, Choi SY, Park J, Lee BH, et al. Nuclear localization and functional characteristics of voltage-gated potassium channel Kv1. 3. Journal of Biological Chemistry. 2015;290(20):12547–57.

40. Panyi G, Bagdany M, Bodnar A, Vamosi G, Szentesi G, Jenei A, et al. Colocalization and nonrandom distribution of Kv1. 3 potassium channels and CD3 molecules in the plasma membrane of human T lymphocytes. Proceedings of the National Academy of Sciences. 2003;100(5):2592–7.

41. Nicolaou SA, Neumeier L, Steckly A, Kucher V, Takimoto K, Conforti L. Localization of Kv1. 3 channels in the immunological synapse modulates the calcium response to antigen stimulation in T lymphocytes. The Journal of Immunology. 2009;183(10):6296–302.

42. Cho KF, Branon TC, Udeshi ND, Myers SA, Carr SA, Ting AY. Proximity labeling in mammalian cells with TurboID and split-TurboID. Nature Protocols. 2020;15(12):3971–99.

43. Valencia-Cruz G, Shabala L, Delgado-Enciso I, Shabala S, Bonales-Alatorre E, Pottosin II, et al. Kbg and Kv1. 3 channels mediate potassium efflux in the early phase of apoptosis in Jurkat T lymphocytes. American Journal of Physiology-Cell Physiology. 2009;297(6):C1544–C53.

44. Doczi MA, Damon DH, Morielli AD. A C-terminal PDZ binding domain modulates the function and localization of Kv1. 3 channels. Experimental cell research. 2011;317(16):2333–41.

45. Rayaprolu S, Bitarafan S, Santiago JV, Betarbet R, Sunna S, Cheng L, et al. Cell type-specific biotin labeling in vivo resolves regional neuronal and astrocyte proteomic differences in mouse brain. Nature Communications. 2022;13(1):2927.

46. Sunna S, Bowen C, Zeng H, Rayaprolu S, Kumar P, Bagchi P, et al. Cellular proteomic profiling using proximity labeling by TurboID-NES in microglial and neuronal cell lines. Molecular & Cellular Proteomics. 2023;22(6).

47. Perez-Riverol Y, Bai J, Bandla C, Garcia-Seisdedos D, Hewapathirana S, Kamatchinathan S, et al. The PRIDE database resources in 2022: a hub for mass spectrometry-based proteomics evidences. Nucleic Acids Res. 2022;50(D1):D543–D52.

48. Szklarczyk D, Gable AL, Lyon D, Junge A, Wyder S, Huerta-Cepas J, et al. STRING v11: protein– protein association networks with increased coverage, supporting functional discovery in genome-wide experimental datasets. Nucleic acids research. 2019;47(D1):D607–D13.

49. Csépányi-Kömi R, Sáfár D, Grósz V, Tarján ZL, Ligeti E. In silico tissue-distribution of human Rho family GTPase activating proteins. Small GTPases. 2013;4(2):90–101.

50. Kelber JA, Klemke RL. PEAK1, a novel kinase target in the fight against cancer. Oncotarget. 2010;1(3):219.

51. Taylor GS, Maehama T, Dixon JE. Myotubularin, a protein tyrosine phosphatase mutated in myotubular myopathy, dephosphorylates the lipid second messenger, phosphatidylinositol 3-phosphate. Proc Natl Acad Sci U S A. 2000;97(16):8910–5.

52. Singla A, Boesch DJ, Fung HYJ, Ngoka C, Enriquez AS, Song R, et al. Structural basis for Retriever-SNX17 assembly and endosomal sorting. Nat Commun. 2024;15(1):10193.

53. Marks DR, Fadool DA. Post-synaptic density perturbs insulin-induced Kv1.3 channel modulation via a clustering mechanism involving the SH3 domain. J Neurochem. 2007;103(4):1608–27.

54. Szilagyi O, Boratko A, Panyi G, Hajdu P. The role of PSD-95 in the rearrangement of Kv1.3 channels to the immunological synapse. Pflugers Arch. 2013;465(9):1341–53.

55. Zeng J, Li M, Shi H, Guo J. Upregulation of FGD6 Predicts Poor Prognosis in Gastric Cancer. Front Med (Lausanne). 2021;8:672595.

56. Hu Q, Guo C, Li Y, Aronow BJ, Zhang J. LMO7 mediates cell-specific activation of the Rho-myocardin-related transcription factor-serum response factor pathway and plays an important role in breast cancer cell migration. Mol Cell Biol. 2011;31(16):3223–40.

57. Amano M, Nakayama M, Kaibuchi K. Rho-kinase/ROCK: A key regulator of the cytoskeleton and cell polarity. Cytoskeleton (Hoboken). 2010;67(9):545–54.

58. Fan L, Tang Y, Li J, Huang W. Increased expression of TBC1D10B as a potential prognostic and immunotherapy relevant biomarker in liver hepatocellular carcinoma. Sci Rep. 2023;13(1):335.

59. Maurin J, Morel A, Guerit D, Cau J, Urbach S, Blangy A, et al. The Beta-Tubulin Isotype TUBB6 Controls Microtubule and Actin Dynamics in Osteoclasts. Front Cell Dev Biol. 2021;9:778887.

60. Levite M, Cahalon L, Peretz A, Hershkoviz R, Sobko A, Ariel A, et al. Extracellular K(+) and opening of voltage-gated potassium channels activate T cell integrin function: physical and functional association between Kv1.3 channels and beta1 integrins. J Exp Med. 2000;191(7):1167–76.

61. Pattu V, Qu B, Schwarz EC, Strauss B, Weins L, Bhat SS, et al. SNARE protein expression and localization in human cytotoxic T lymphocytes. Eur J Immunol. 2012;42(2):470–5.

62. Desvignes L, Weidinger C, Shaw P, Vaeth M, Ribierre T, Liu M, et al. STIM1 controls T cell-mediated immune regulation and inflammation in chronic infection. J Clin Invest. 2015;125(6):2347–62.

63. Ngoenkam J, Schamel WW, Pongcharoen S. Selected signalling proteins recruited to the T-cell receptor-CD3 complex. Immunology. 2018;153(1):42–50.

64. Panyi G, Vamosi G, Bacso Z, Bagdany M, Bodnar A, Varga Z, et al. Kv1.3 potassium channels are localized in the immunological synapse formed between cytotoxic and target cells. Proc Natl Acad Sci U S A. 2004;101(5):1285–90.

65. Canas CA, Castano-Valencia S, Castro-Herrera F. Pharmacological blockade of KV1.3 channel as a promising treatment in autoimmune diseases. J Transl Autoimmun. 2022;5:100146.

66. Fung-Leung WP, Edwards W, Liu Y, Ngo K, Angsana J, Castro G, et al. T Cell Subset and Stimulation Strength-Dependent Modulation of T Cell Activation by Kv1.3 Blockers. PLoS One. 2017;12(1):e0170102.

67. Chandy KG, Norton RS. Peptide blockers of K(v)1.3 channels in T cells as therapeutics for autoimmune disease. Curr Opin Chem Biol. 2017;38:97–107.

68. Zhang R, Kang R, Tang D. The STING1 network regulates autophagy and cell death. Signal Transduction and Targeted Therapy. 2021;6(1):208.

69. Sebestyen V, Nagy E, Mocsar G, Volko J, Szilagyi O, Kenesei A, et al. Role of C-Terminal Domain and Membrane Potential in the Mobility of Kv1.3 Channels in Immune Synapse Forming T Cells. Int J Mol Sci. 2022;23(6).

70. Butte MJ, Stein JV, Delon J. Editorial: The cytoskeleton in T cell migration and activation. Frontiers in Immunology. 2022;13.

71. Saoudi A, Kassem S, Dejean AS, Gaud G. Rho-GTPases as key regulators of T lymphocyte biology. Small GTPases. 2014;5(4):e983862.

72. Huang J, Dibble CC, Matsuzaki M, Manning BD. The TSC1-TSC2 complex is required for proper activation of mTOR complex 2. Mol Cell Biol. 2008;28(12):4104–15.

73. Xu J, Koni PA, Wang P, Li G, Kaczmarek L, Wu Y, et al. The voltage-gated potassium channel Kv1.3 regulates energy homeostasis and body weight. Human Molecular Genetics. 2003;12(5):551–9.

74. Zhao L, Gao N, Peng X, Chen L, Meng T, Jiang C, et al. TRAF4-Mediated LAMTOR1 Ubiquitination Promotes mTORC1 Activation and Inhibits the Inflammation-Induced Colorectal Cancer Progression. Adv Sci (Weinh). 2024;11(12):e2301164.

75. Nalkurthi C, Schroder WA, Melino M, Irvine KM, Nyuydzefe M, Chen W, et al. ROCK2 inhibition attenuates profibrogenic immune cell function to reverse thioacetamide-induced liver fibrosis. JHEP Rep. 2022;4(1):100386.

76. Jiang L, Zhu X, Yang H, Chen T, Lv K. Bioinformatics Analysis Discovers Microtubular Tubulin Beta 6 Class V (TUBB6) as a Potential Therapeutic Target in Glioblastoma. Front Genet. 2020;11:566579.

77. Nygaard HB, van Dyck CH, Strittmatter SM. Fyn kinase inhibition as a novel therapy for Alzheimer’s disease. Alzheimers Res Ther. 2014;6(1):8.

78. Angelopoulou E, Paudel YN, Julian T, Shaikh MF, Piperi C. Pivotal Role of Fyn Kinase in Parkinson’s Disease and Levodopa-Induced Dyskinesia: a Novel Therapeutic Target? Mol Neurobiol. 2021;58(4):1372–91.

79. Sarkar S, Nguyen HM, Malovic E, Luo J, Langley M, Palanisamy BN, et al. Kv1.3 modulates neuroinflammation and neurodegeneration in Parkinson’s disease. J Clin Invest. 2020;130(8):4195–212.

80. Rangaraju S, Gearing M, Jin LW, Levey A. Potassium channel Kv1.3 is highly expressed by microglia in human Alzheimer’s disease. J Alzheimers Dis. 2015;44(3):797–808.

81. Ye W, Zhou Y, Xu B, Zhu D, Rui X, Xu M, et al. CD247 expression is associated with differentiation and classification in ovarian cancer. Medicine (Baltimore). 2019;98(51):e18407.

82. Chinen T, Kannan AK, Levine AG, Fan X, Klein U, Zheng Y, et al. An essential role for the IL-2 receptor in Treg cell function. Nature Immunology. 2016;17(11):1322–33.

83. Hu B, Duan S, Wang Z, Li X, Zhou Y, Zhang X, et al. Insights Into the Role of CSF1R in the Central Nervous System and Neurological Disorders. Front Aging Neurosci. 2021;13:789834.

84. Ponce LP, Fenn NC, Moritz N, Krupka C, Kozik JH, Lauber K, et al. SIRPalpha-antibody fusion proteins stimulate phagocytosis and promote elimination of acute myeloid leukemia cells. Oncotarget. 2017;8(7):11284–301.

85. Gaultier A, Wu X, Le Moan N, Takimoto S, Mukandala G, Akassoglou K, et al. Low-density lipoprotein receptor-related protein 1 is an essential receptor for myelin phagocytosis. J Cell Sci. 2009;122(Pt 8):1155–62.

86. Gulbins E, Sassi N, Grassme H, Zoratti M, Szabo I. Role of Kv1.3 mitochondrial potassium channel in apoptotic signalling in lymphocytes. Biochim Biophys Acta. 2010;1797(6-7):1251–9.

87. Reyes A, Melchionda L, Burlina A, Robinson AJ, Ghezzi D, Zeviani M. Mutations in TIMM50 compromise cell survival in OxPhos-dependent metabolic conditions. EMBO Mol Med. 2018;10(10).

88. Dudek J, Rehling P, van der Laan M. Mitochondrial protein import: common principles and physiological networks. Biochim Biophys Acta. 2013;1833(2):274–85.

89. Ng MYW, Wai T, Simonsen A. Quality control of the mitochondrion. Dev Cell. 2021;56(7):881–905.

90. Zhang L, Li M, Chan AK, Liu Q, Kang H, Pokharel SP, et al. COX4I1 Controls Mitochondrial Electron Transport Chain Complex IV Assembly and Leukemia Progression in Acute Myeloid Leukemia. Blood. 2023;142(Supplement 1):581-.

91. Ohsakaya S, Fujikawa M, Hisabori T, Yoshida M. Knockdown of DAPIT (diabetes-associated protein in insulin-sensitive tissue) results in loss of ATP synthase in mitochondria. J Biol Chem. 2011;286(23):20292–6.

92. Unni S, Thiyagarajan S, Srinivas Bharath MM, Padmanabhan B. Tryptophan Oxidation in the UQCRC1 Subunit of Mitochondrial Complex III (Ubiquinol-Cytochrome C Reductase) in a Mouse Model of Myodegeneration Causes Large Structural Changes in the Complex: A Molecular Dynamics Simulation Study. Sci Rep. 2019;9(1):10694.

93. Fairley LH, Wong JH, Barron AM. Mitochondrial Regulation of Microglial Immunometabolism in Alzheimer’s Disease. Front Immunol. 2021;12:624538.

94. Steinert EM, Vasan K, Chandel NS. Mitochondrial Metabolism Regulation of T Cell-Mediated Immunity. Annu Rev Immunol. 2021;39:395–416.

95. Nguyen W, Howard BL, Neale DS, Thompson PE, White PJ, Wulff H, et al. Use of Kv1.3 blockers for inflammatory skin conditions. Curr Med Chem. 2010;17(26):2882–96.

96. Yuan XL, Zhao YP, Huang J, Liu JC, Mao WQ, Yin J, et al. A Kv1.3 channel-specific blocker alleviates neurological impairment through inhibiting T-cell activation in experimental autoimmune encephalomyelitis. CNS Neurosci Ther. 2018;24(10):967–77.

97. Lintermans LL, Stegeman CA, Munoz-Elias EJ, Tarcha EJ, Iadonato SP, Rutgers A, et al. Kv1.3 blockade by ShK186 modulates CD4+ effector memory T-cell activity of patients with granulomatosis with polyangiitis. Rheumatology (Oxford). 2024;63(1):198–208.

98. Khodoun M, Chimote AA, Ilyas FZ, Duncan HJ, Moncrieffe H, Kant KS, et al. Targeted knockdown of Kv1.3 channels in T lymphocytes corrects the disease manifestations associated with systemic lupus erythematosus. Sci Adv. 2020;6(47).

99. Lioudyno MI, Kozak JA, Penna A, Safrina O, Zhang SL, Sen D, et al. Orai1 and STIM1 move to the immunological synapse and are up-regulated during T cell activation. Proc Natl Acad Sci U S A. 2008;105(6):2011–6.

100. Cahalan MD, Chandy KG. The functional network of ion channels in T lymphocytes. Immunol Rev. 2009;231(1):59–87.

