## Supplementary Figures for "Identification of novel Kv1.3 channel-interacting proteins using proximity labelling in T-cells"

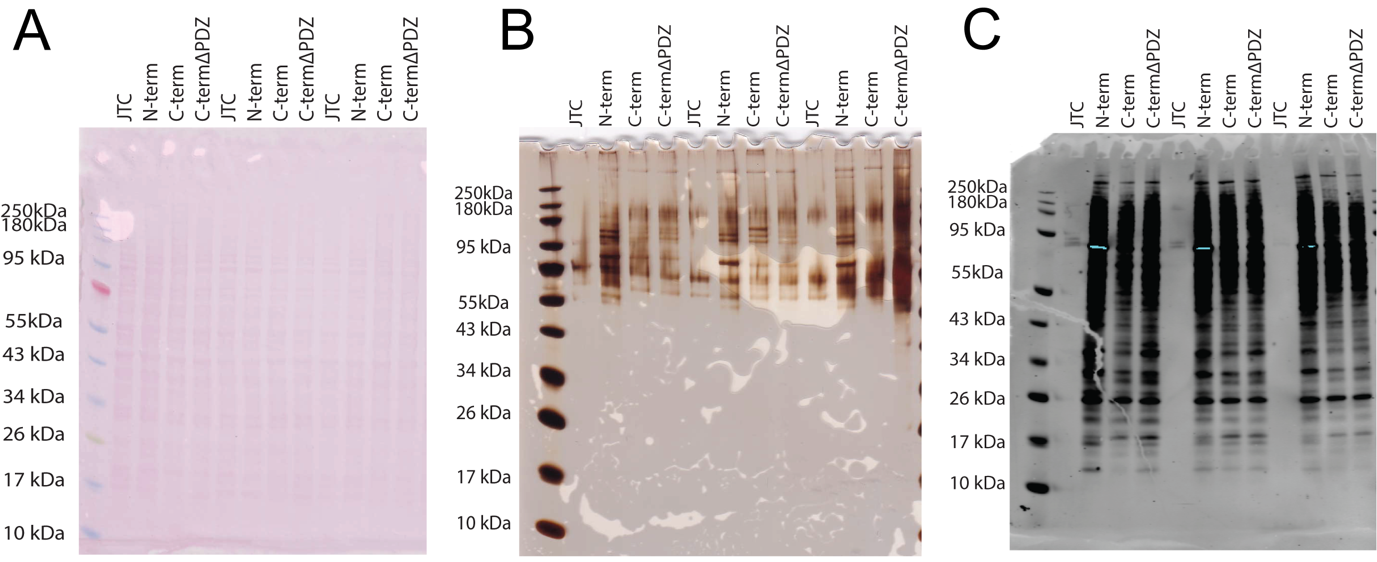

**Supplemental Figure 1:** **Protein integrity of Kv1.3-TurboID transduced Jurkat T-cells maintained throughout processing of lysates.** **(A)** Ponceau staining of protein prior to probing for western blot highlights even protein loading and distribution (n=3). **(B)** Post-Affinity purification silver stain shows successful purification of proteins present after AP (n=3). **(C)** Post-affinity purification western blot utilizing streptavidin-680 shows enrichment of biotinylated proteins following AP (n=3).

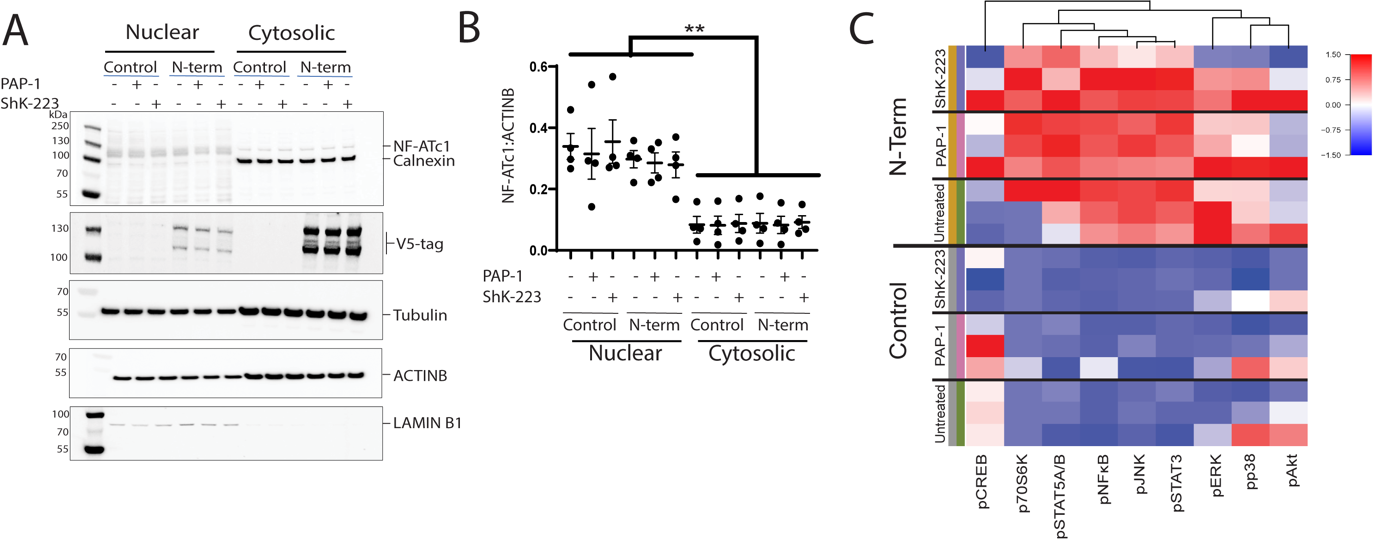

**Supplemental Figure 2: Kv1.3 channel transduction effects on Jurkat T-cells properties. (A)** Western blot depicting effect of Kv1.3 channel transduction and blockade on NF-AT signaling (n=4). **(B)** Densitometric analysis of western blot given in Fig 2A. Band density was analyzed in ImageJ software (n=4). **(C)** Heat map to represent effect of Kv1.3-TurboID constructs transduction in Jurkat T-cells and channel blockade by PAP-1 (100 nM) and ShK-223 (200 nM) on multiple signaling proteins measured by multiplex assay (n=3). Statistical analyses were done by One-way ANOVA analysis, followed by Tukey’s test. p < 0.01 (∗∗). Statistical analysis of Luminex data is given in Supp. Datasheet 2.

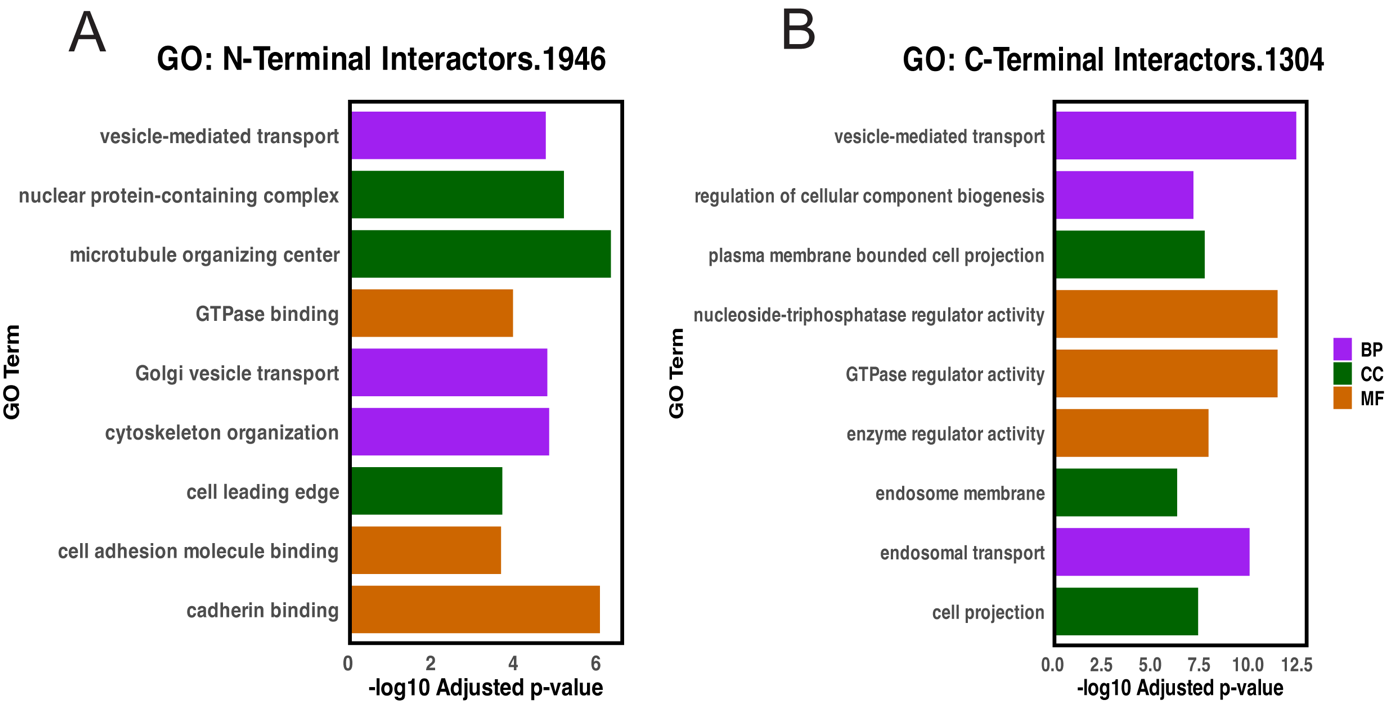

**Supplemental Figure 3: Pathway enrichment analysis of Kv1.3 terminal specific interactors. (A)** Gene Ontology (GO) term enrichment analysis of N-terminal interactors identified in Fig 3A shows Kv1.3 channel’s N-terminal role in protein trafficking and localization. **(B)** GO analysis of C-terminal interactors identified in Fig 3B displays C-terminal functioning in protein transport and regulation of GTPase activity. All related analyses are provided in Supp. Datasheet 4.

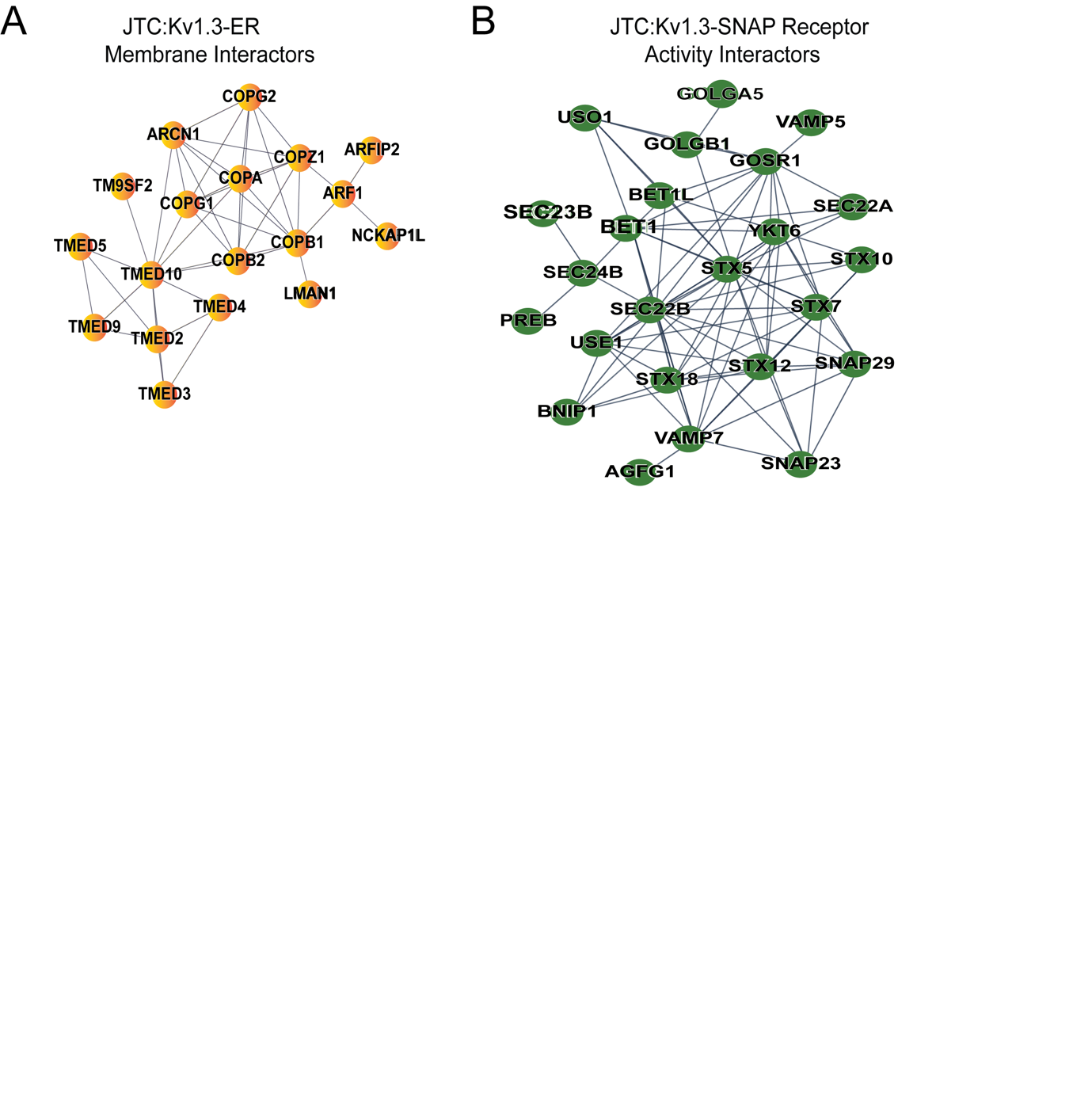

**Supplemental Figure 4: Network analysis of membrane interactors of Kv1.3 channel in Jurkat T-cells. (A)** STRING analysis of 335 membrane interactors of Kv1.3 channel identified in Fig 5A shows 18 ER membrane proteins interacting with Kv1.3 channel in JTCs. **(B)** Network analysis of Kv1.3 channel’s membrane interactors in JTCs highlights 25 proteins from SNAP receptor activity family. Details of related analyses are provided in Supp. Datasheet 6.
